# Integrin diversity enables dynamic adhesion control in a homeostatic epithelium

**DOI:** 10.64898/2026.08.13.744583

**Authors:** Jia Chen, Daiki Sugita, Edward Allgeyer, Dhriti Saumya, DhanLakshmi Shunmugam, Daniel St Johnston

## Abstract

Homeostatic epithelia must balance stem cell maintenance, progenitor differentiation, and clearance of damaged cells while preserving barrier integrity. We investigated how integrin– ECM adhesion is regulated in the *Drosophila* midgut, a homeostatic epithelium with basal stem cells. The midgut expresses two beta integrins: ubiquitous βMys and endoderm-specific βν. ISCs and enteroblasts express only βMys, which pairs with αMew to mediate enteroblast attachment to the basement membrane. In contrast, enterocytes express both βMys and βν; αMew/βν supports ECM adhesion, while βMys pairs with αScab and localises to the basal labyrinth. Enterocytes lacking αMew or βν detach and are apically extruded, but this phenotype is rescued when the corresponding integrin is removed from the entire epithelium. Thus, enterocytes compete for basement membrane adhesion, with less adhesive cells being eliminated by their neighbours. In βν homozygotes, enteroblasts expand basally and adopt a migratory-like morphology. We propose that integrin-mediated competition for ECM adhesion is a general phenomenon that functions in the midgut to promote enterocyte extrusion, which stimulates the migration of nearby enteroblasts to maintain gut homeostasis.

## Introduction

Cell-extracellular matrix (ECM) adhesion is fundamental to epithelial biology, underpinning cell migration, mechanosensing, and the establishment of apical-basal polarity. Epithelial cells engage the ECM primarily through integrins, which function as αβ heterodimers that are primed, activated, and clustered at the cell-ECM interface. The extracellular domain of the α subunit confers ligand specificity for ECM components such as laminins or collagens, while the β subunit’s cytoplasmic tail recruits intracellular proteins like Talin and Kindlin [1]. These interactions trigger conformational changes that activate integrin function and link adhesions to the cytoskeleton. Integrin-mediated adhesions support both dynamic focal contacts in migrating cells and stable basal interactions in steady-state epithelia [2]. While integrin clustering is well established for mechanosensing and turnover during migration, far less is known about how epithelia manage multiple co-expressed αβ heterodimers under homeostatic conditions to maintain tissue integrity, polarity, and stem cell regulation.

Unlike vertebrates, which encode over 20 integrin subunits to generate diverse heterodimers, *Drosophila melanogaster* expresses only five α and two β subunits Myospheroid (βMys, also known as βPS integrin) and βν [3]. This simplicity makes flies an ideal model to dissect subunit-specific functions. βMys is expressed and required in multiple tissues during development, while βν is only expressed in endoderm-derived tissues such as the midgut and appears dispensable: *βν* mutants are viable and fertile [4].

The adult fly midgut consists of a simple layer of epithelial cells, with their apical surfaces facing the gut lumen and their basal sides contacting the basement membrane, which lies immediately above the visceral muscles, trachea and nerves. Enterocyte apical–basal polarity differs from other fly epithelia, as it does not depend on canonical polarity factors but instead relies on basal signals [5]. The midgut epithelium arises from the embryonic endoderm, where βMys is the primary β integrin that drives migration [4,6]. In adult flies, the epithelium is homeostatic: basally located intestinal stem cells (ISCs) divide asymmetrically to produce post-mitotic enteroblasts (EBs) that differentiate into enterocytes as they insert into the epithelium to replace dying cells [7,8]. βMys is important for ISC maintenance, while βν is not essential for either embryonic midgut morphogenesis or adult ISC proliferation [4,9–11]. Basal integrins in the ECs anchor the epithelium to the basement membrane and suppress precocious ISC activity [12–14]. Despite these findings, integrin functions across the ISC/EB-EC lineage, and their precise contributions to enterocyte adhesion remain undefined.

A key unresolved question is why the midgut epithelium, with its unique epithelial cell polarity, resident stem cells, and dynamic cell turnover, requires a second β integrin. Here, we map the localization of integrin subunits with high-resolution across the ISC/EB-EC lineage and uncover distinct, non-redundant functions of βMys and βν containing Integrin dimers. Specifically, we reveal their cooperative and competitive contributions to cell type-specific basal adhesion, EB reattachment post-division, and epithelial homeostasis.

## Results

### βν is not localized or required in ISC/early EBs

Early studies using RT-PCR on the fly midgut revealed that four out of the five alpha Integrin subunits: *α1*-*multiple edematous wings* (*αmew*), *α2*-*inflated* (*αif*), *α-scab* (*α4)* and *a4*, and both beta Integrins: *βmys* and *βν*, are expressed in the midgut[9]. Dissected fly midgut tissue contains muscles, trachea, and neurons as well as the gut epithelium, which contains enterocytes and enteroendocrine cells, their precursors and intestinal stem cells (ISCs). Since 90% of the epithelial cells are enterocytes (ECs), we first examined integrin expression and localization in ECs and their precursors, the enteroblasts (EBs).

The Fly Cell Atlas shows high expression of both beta integrins (βMys, βν) and two alpha subunits (αMew, αScab) in ECs, with αIf mainly in muscle and α4 barely detectable. In ISC/EB clusters, βν exhibits the lowest enrichment compared to αMew, αScab, and βMys[15]. Because of the lack of tools (transgenic fly lines or reagents) targeting α4, we visualized the cellular localization of the four other integrins, αMew, αScab, βMys and βν in ISC/EBs and ECs using antibody staining and functional protein trap lines (for βν, we used both antibody staining and a strain expressing a functional GFP-tagged transgene under the control of its own promoter [16], Figure 1A and Figure S1A). Consistent with the Fly cell Atlas findings and previous studies [9,15], αMew and βν are both highly expressed in enterocytes and localize to the basal domain (Figure 1A and Figure S1A). Although they may also be weakly expressed in ISC/EBs, it is hard to distinguish between signal from out of focus ECs and the ISC/EBs, due the latter’s small size and scattered distribution in the steady-state epithelium (Figure 1A, yellow arrow). To analyse integrin subunit expression and localization specifically in ISC/EBs, we therefore used GFP/YFP-RNAi lines driven by *MyoIA^ts^* to suppress the signal in ECs and unmask integrin patterns in these smaller cells.

**Figure 1.**
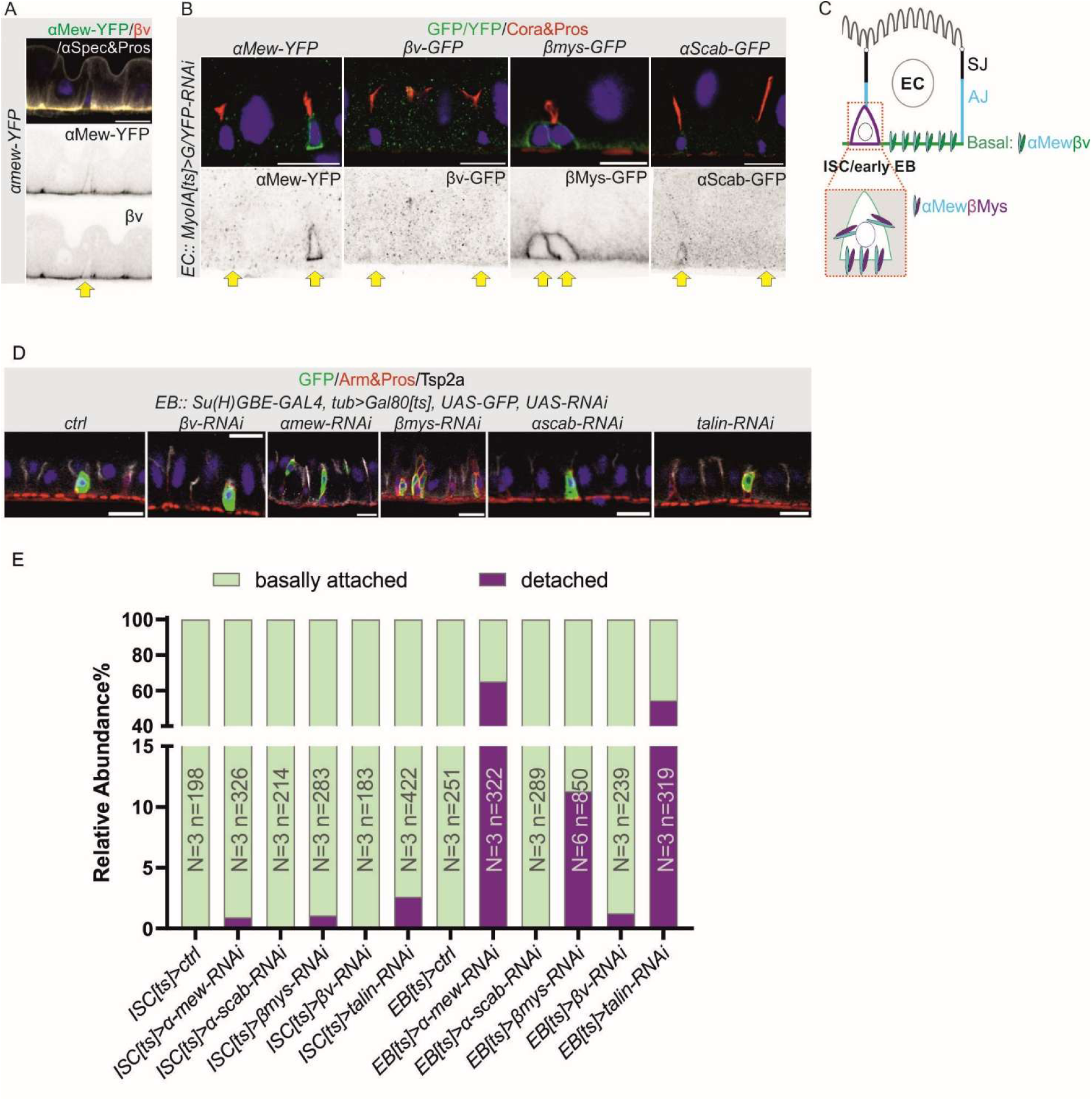
βν is not required or localized at the cell membrane in ISC/early EBs. A. A side view of a region of the midgut epithelium expressing *αmew-YFP* and stained for YFP (green), βν (red) and αSpectrin and Pros (grayscale; αSpectrin labels the cell membrane and Pros labels enteroendocrine (ee) cells). Single channels of αMew-YFP and βν are shown below in inverted grayscale. In enterocytes, both αMew-YFP and βν localize to the basal domain, while βν is barely detectable in the ISC/EB (yellow arrow). B. A side view of the midgut epithelium showing the expression and localisation of *αmew-YFP*, *βmys-GFP* and *αscab-GFP* protein trap lines and a *βν-GFP* genomic transgene in ISCs/EBs (GFP/YFP, green; Cora (Septate Junctions (SJ)) and Pros (ee), red). The expression of the GFP-tagged integrins was knocked down in ECs using the enterocyte-specific driver, *MyoIA*-Gal4, to express UAS-GFP RNAi. Individual GFP or YFP channels are shown below in inverted grayscale. Yellow arrows indicate the ISCs/EBs. C. A diagram illustrating the localisation of the different integrin subunits in enterocytes (EC) and ISC/early EBs. αMew and βν localize to the basal domain in ECs, while αMew and Mys localize around the plasma membrane of ISC/EBs. Septate Junction (SJ), Adherens Junction (AJ). D. A side view of the midgut epithelium after knocking down individual integrin subunits and Talin specifically in enteroblasts using *Su(H)GBE-GAL4; tub>GAL80[ts], UAS-GFP, UAS-RNAi*, showing GFP+ EBs in green, Arm (AJ) & Pros (ee) in red and Tsp2a (SJ) in grayscale. E. Quantification of the relative abundance of basally-attached (green) or detached (purple) ISCs/EBs. The slight increase of detaching cells in *ISC*^ts^>*αmew/βmys-RNAi* can be explained by the newborn enteroblast is most likely detached from the BM after asymmetric cell division. Knocking down Talin, the main intracellular Integrin scaffolding protein and activator recapitulates the effects of knocking down integrins in EBs. The small number of detaching cells in *ISC*^ts^>*αmew/βmys/talin-RNAi* may be due to newborn daughter cell detaching after the ISC has asymmetrically divided. N, total number of experiments; n, total number of cells. Scale bars,10μm.

**Figure S1.**
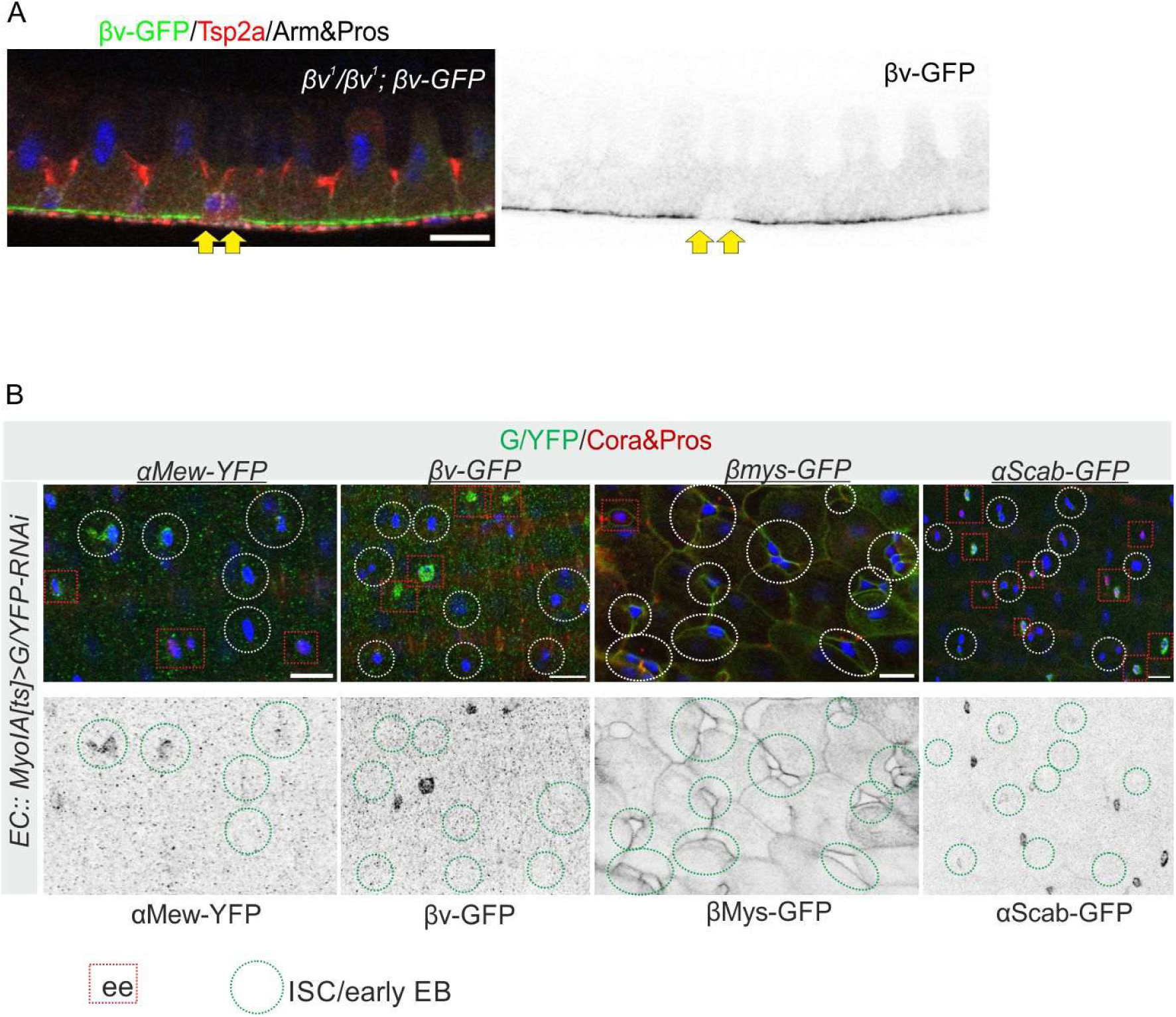
βν is not required or localized at the cell membrane in ISC/early EBs. A. Immunostaining for GFP (*βν*, green), Tsp2a (SJ, red), Arm (AJ) and Pros (grayscale) in *βν*^1^/ *βν*^1^; *βν-GFP*. βν localizes to the basal domain in enterocytes but is absent in ISC/early EBs (yellow arrows). B. A basal view of the midgut epithelium as in Figure 1B. Red squares highlight enteroendocrine cells (ee) and green circles highlight ISCs/early EBs. βν localizes to the basal domain in ees. Scale bars, 10μm.

Under steady-state conditions without transcriptional markers, ISCs and early EBs are morphologically identical until the EBs start to differentiate and endo-replicate; we therefore classified Pros⁻ small cells as ISC/early EBs. Surprisingly, βν was absent from these cells, whereas αMew and βMys showed strong membrane localization (basal/lateral), while αScab was weakly expressed (Figure 1B and Figure S1B). Anti-βν staining confirmed its poor expression or localization in ISC/EBs (Figures 1A and S1A). These findings suggest βMys pairs mainly with αMew in ISC/early EBs and localizes basally and laterally, while βν pairs with αMew in ECs and localises to the basal domain. (Figure 1C).

We tested the function of Integrins in ISCs or enteroblasts by depleting each subunit using cell-type specific RNAi. Consistent with several previous studies [9,10], we found that knock down of all individual integrin subunits (including βν) had no effect on ISC attachment to the basement membrane. Similarly, loss of βν in EBs had no phenotype in terms of cell attachment or shape. In contrast, losing either αMew or βMys in EBs caused basal detachment, creating small cells floating within the epithelium (Figure 1D and 1E). Thus, βν is neither localized nor required in ISC/EBs, whereas αMew and βMys localize to the plasma membrane and are necessary for enteroblast attachment.

### αScab/Mys and αMew/βν localize to different membrane domains in enterocytes

To determine the relationships between the two alpha integrins (αMew and αScab) and two beta integrins (βMys and βν) in the enterocytes, we imaged endogenously-tagged versions of each protein. As already shown in Figure 1A, αMew and βν co-localise at the basal domain of the ECs. In *αmew-YFP, βmys-mCherry* midguts, βMys localises to basal membrane infoldings called the basal labyrinth (BL), whereas αMew occupies the basal domain from which the basal infoldings arise (Figure 2A1 and Figure S2C). αScab also localizes to basal labyrinth in enterocytes (Figure 2C, Figure S2A and S2B). This suggests that αScab pairs with βMys, while αMew dimerises with βν in ECs. It is worth noting that the basal domain and basal labyrinth form a continuous membrane structure, but the ECM only underlies the basal domain and is not visible in the lumen of the basal labyrinth by electron microscopy [17].

**Figure 2.**
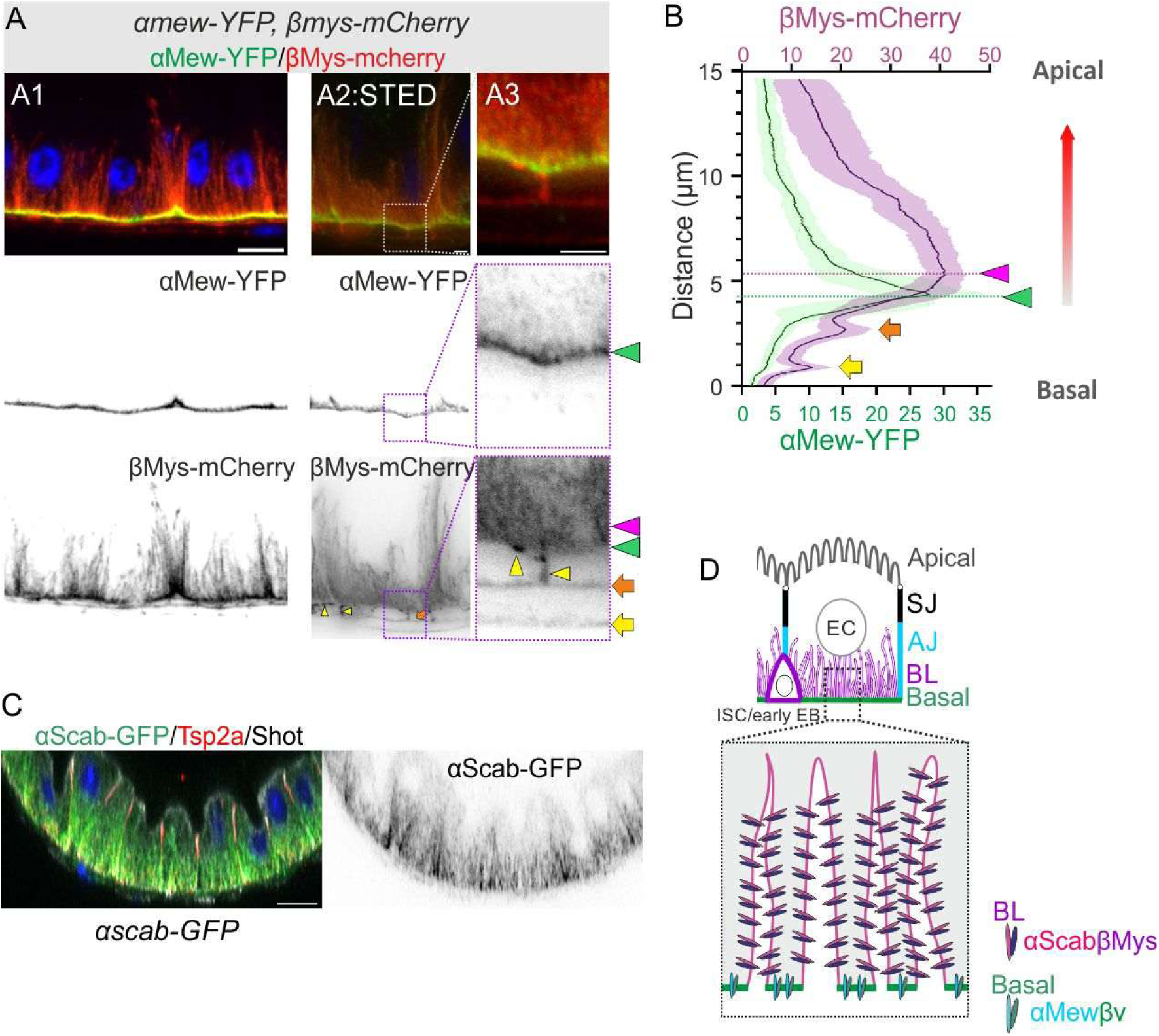
αScab/βMys and αMew/βν localize to different membrane domains in enterocytes. A. A side view of an *αmew-YFP, βmys-mCherry* midgut epithelium showing that αMew-YFP (green) localises to the basal domain and βMys-mcherry (red) to the basal labyrinth. Separate channels of the αMew-YFP and βMys-mCherry are shown below in inverted grayscale. A1: images obtained on a confocal microscope, scale bar, 10μm; A2 and A3 (magnified image of A2): images obtained with a STED microscope, scale bar, 2μm. βMys is also expressed in the visceral muscles. The yellow arrowheads indicate βMys in the circular muscle layer, the orange arrow indicates the boundary between the circular and longitudinal muscle and the yellow arrow marks the bottom of the longitudinal muscle layer. In the muscles, βMys presumably forms dimers with *αinflated*, which shows a very similar localisation (Figure S2D). B. Quantification of the average fluorescence intensity along the enterocyte apical to basal axis of αMew-YFP (green) and βMys-mCherry (purple) in STED images (3 Cells; 5 measurements each). Note that βMys intensity peaks inside the ECs (purple arrowhead) above the αMew intensity peak (green arrowhead) at the basal domain. βMys also localises around the longitudinal muscle layer (orange and yellow arrows). C. A side view of an *αscab-GFP* midgut epithelium, showing αScab (green) localization to the basal labyrinth, Tsp2a (SJ; red) and Shot (apical; grayscale). The αScab-GFP channel is shown in inverted grayscale on the right. Scale bar=10μm. D. An illustration depicting the localisation of αScab/βMys to the basal labyrinth (BL) and αMew/βν to the basal domain of the enterocytes (EC).

**Figure S2.**
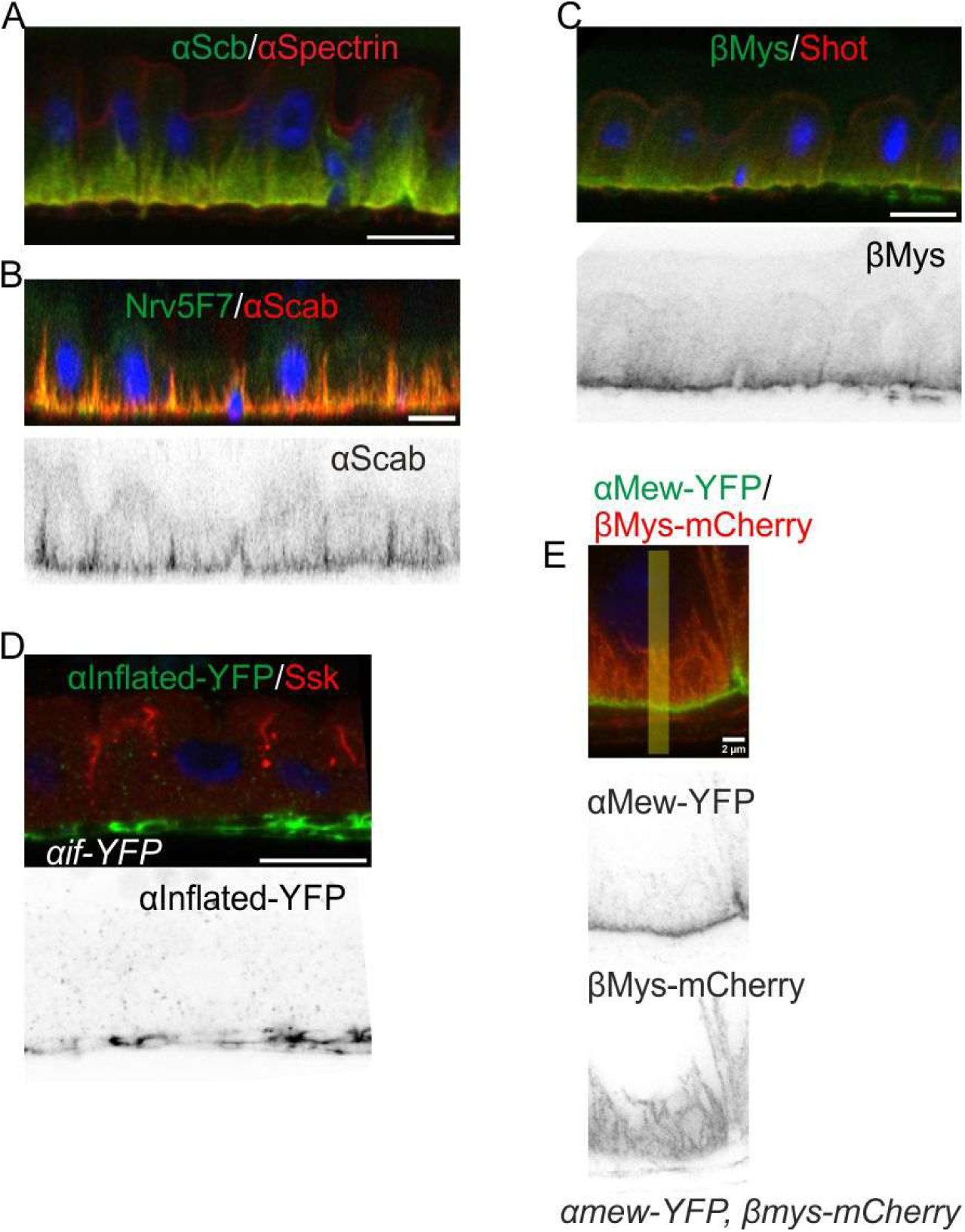
αScab/βMys and αMew/βν localize to different membrane domains in enterocytes. A. A side view of wild-type midgut epithelium stained for Scab (green) and α-Spectrin (red). Both proteins localise to the basal labyrinth. Scale bar=10μm. B. A side view of a wild-type midgut epithelium stained for Scab (red) and Nervana 1, the beta subunit of the Na^+^/K^+^ ATPase (green), using antibody Nrv5F7. Scale bar=10μm. C. A side view of a wild-type midgut epithelium stained for βMys (green) and Shot (red). Scale bar=10μm. D. A side view of a midgut expressing an *αinflated-YFP* genomic transgene, showing α-Inflated (green) localization in the circumferential and longitudinal muscles. Ssk (SJ) in red. Scale bar=10μm. E. Another example of a STED image of *αmew-YFP, βmys-mCherry* midgut epithelium showing αMew-YFP in green and βMys-mCherry in red. The yellow bar is an example of the apical-basal lines used to measure the fluorescence intensity. Scale bar=2μm.

To better resolve the localisation of αScab/βMys relative to αMew/βν, we used Stimulated Emission Depletion (STED) microscopy to image αMew and βMys. This revealed a clear separation between βMys-mCherry and αMew-YFP, with the peak signal for αMew-YFP >1µm below the peak for βMys-mCherry (Figure 2A2, 2A3 and 2B, Figure S2E). Thus, although the basal membrane and basal labyrinth are continuous, αMew/βν dimers localise to the basal membrane in contact with the ECM, while αScab/βMys dimers reside in the basal labyrinth (Figure 2D).

Since βMys/αScab heterodimers localise to the basal labyrinth, we investigated whether they are required for the formation of this structure. However, the basal labyrinth forms normally in both *βmys* and *αscab* null mutant clones, as shown by α-Spectrin staining, which marks the cortical cytoskeleton around the plasma membrane, and by Nrv1, a Na^+^/K^+^ ATPase localized to the basal labyrinth (Figure S3A and Figure 3C). Furthermore, neither mutant had any visible effect on cell shape or cell attachment to the basal membrane, even though βMys plays a role in ECM attachment in the EBs, from which the enterocytes develop (Figure 3A-D, Figure S3A and S3B). This suggests that the 10% of EBs lacking βMys that detach from the basement membrane, subsequently reattach as they grow and differentiate into ECs.

**Figure 3.**
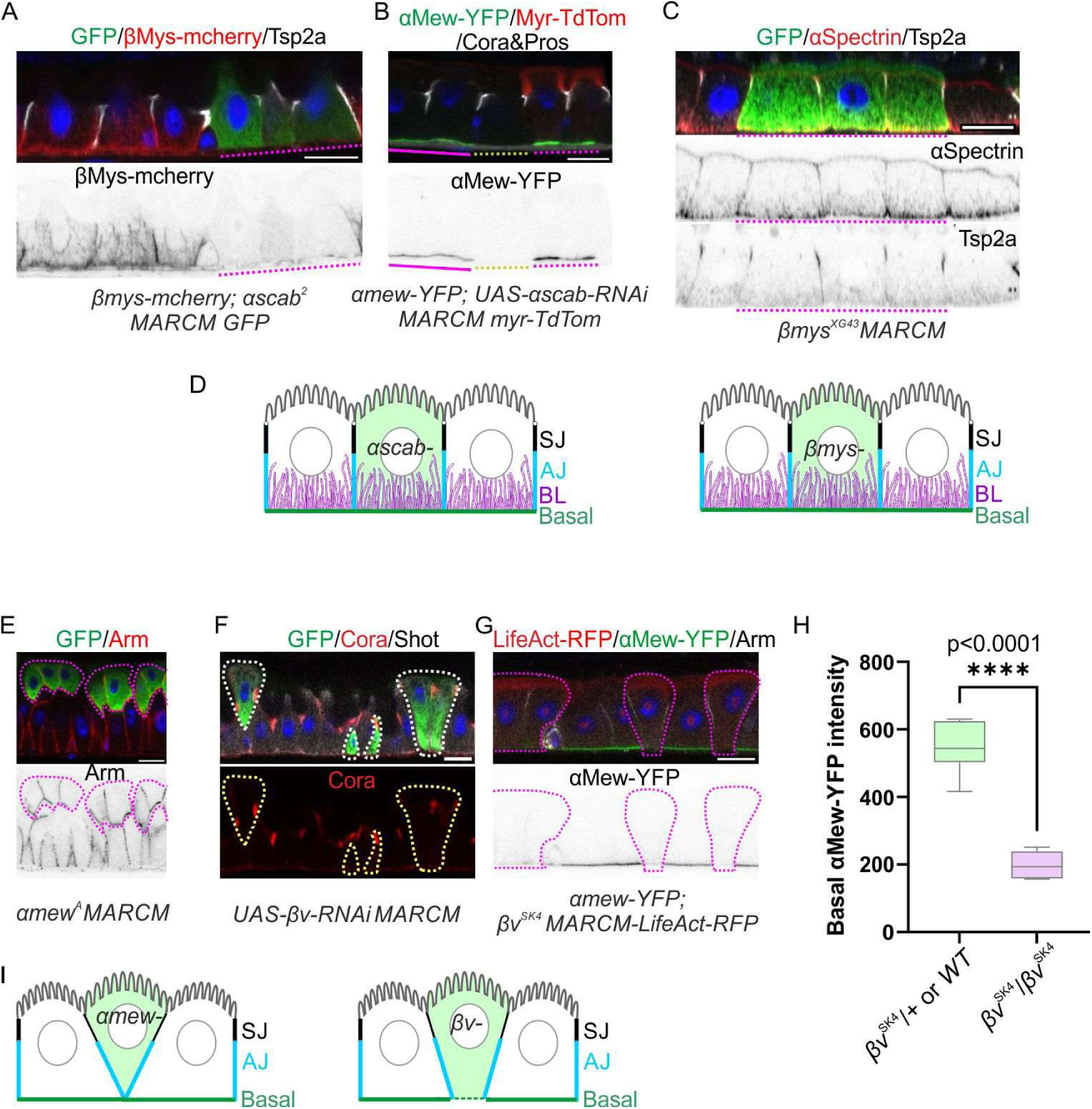
αMew/βν, not αScab/βMys, dimers mediate enterocyte adhesion to the ECM. A. A side view of a midgut containing an *αscab*^2^ MARCM clone (GFP+, green). The *αscab*^2^ mutant cells show the loss of Mys-mCherry (red) in the mutant cells. Tsp2a in grayscale. The Mys-mCherry channel is shown in inverted grayscale below. B. A side view of a midgut containing MARCM clones expressing *UAS-αscab-RNAi*, marked by Myristoylated-TdTomato (Myr-TdTom) + (red). The clones show the same phenotype as *α-scab*^2^ mutant clones, with no defect in basal adhesion. Please note that there are 2 copies of *αmew-YFP* in the clones expressing *UAS-αscab-RNAi*, marked by the expression of Myr-TdTom. The cell in the middle is a twin-spot of the clone and lacks *αmew-YFP*. Single αMew-YFP (green) and Cora(red) channels are shown below in inverted grayscale. The purple dotted line marks the cells expressing *UAS-αscab-RNAi*, the yellow dotted line marks the cell as a twin-spot lacks *αmew-YFP,* and the purple line marks the cells with one copy of *αmew-YFP*. C. A side view of a midgut containing *βmys*^XG43^ MARCM clones (GFP+, green). The mutant cells show no defect in the basal adhesion of the enterocytes or the structure of the basal labyrinth (BL), labeled with αSpectrin (red). Single channels of αSpectrin and Tsp2a are shown below in inverted grayscale. The purple dotted line marks the mutant clones. D. Diagrams show that loss of either βMys or αScab has no effect on the basal adhesion of the enterocytes or the formation of the basal labyrinth. E. A side view of a midgut containing *αmew*^A^ MARCM clones (GFP+, green). Mutant cells are apically extruded into the gut lumen, but form normal adherens junctions, marked by Arm staining (red). The Arm channel is also shown below in inverted grayscale. The purple dotted line marks the mutant clones. F. A side view of MARCM clones (GFP+, green) expressing *UAS-βν-RNAi.* The cells in which *βν* has been knocked down are partially apically extruded into the gut lumen. Cora (SJ) is shown in red and Shot (Apical domain) in grayscale. The dotted line marks the cells expressing *UAS-βν-RNAi*. G. A side view of a midgut containing *βν^SK4^* MARCM clones marked by the expression of LifeActin-RFP (red). Mutant cells are partially extruded apically but maintain some basal contact with the ECM. The clones are outlined with dashed lines. The basal localisation of αMew-YFP (green, and in inverted grayscale below) is strongly reduced in *βν^SK4^* mutant enterocytes. The purple dotted line marks the mutant clones. H. Quantification of the intensity of αMew-YFP fluorescence at the basal domain in *βν^SK4^*clones (purple) compared with the neighboring wild type enterocytes (green). The background fluorescence intensity was deducted from all measurements. These data are collected from 7 guts, with a total of 159 *βν*^SK4^/*βν*^SK4^ mutant cells and 144 neighboring *βν*^SK4^/*+* or wild-type enterocytes. I. Diagram illustrating the apical extrusion of *αmew* mutant enterocytes with little to no basal contact with the ECM, and partial extrusion of *βν* mutant enterocytes with some contact with the ECM. Scale bars, 10μm.

**Figure S3.**
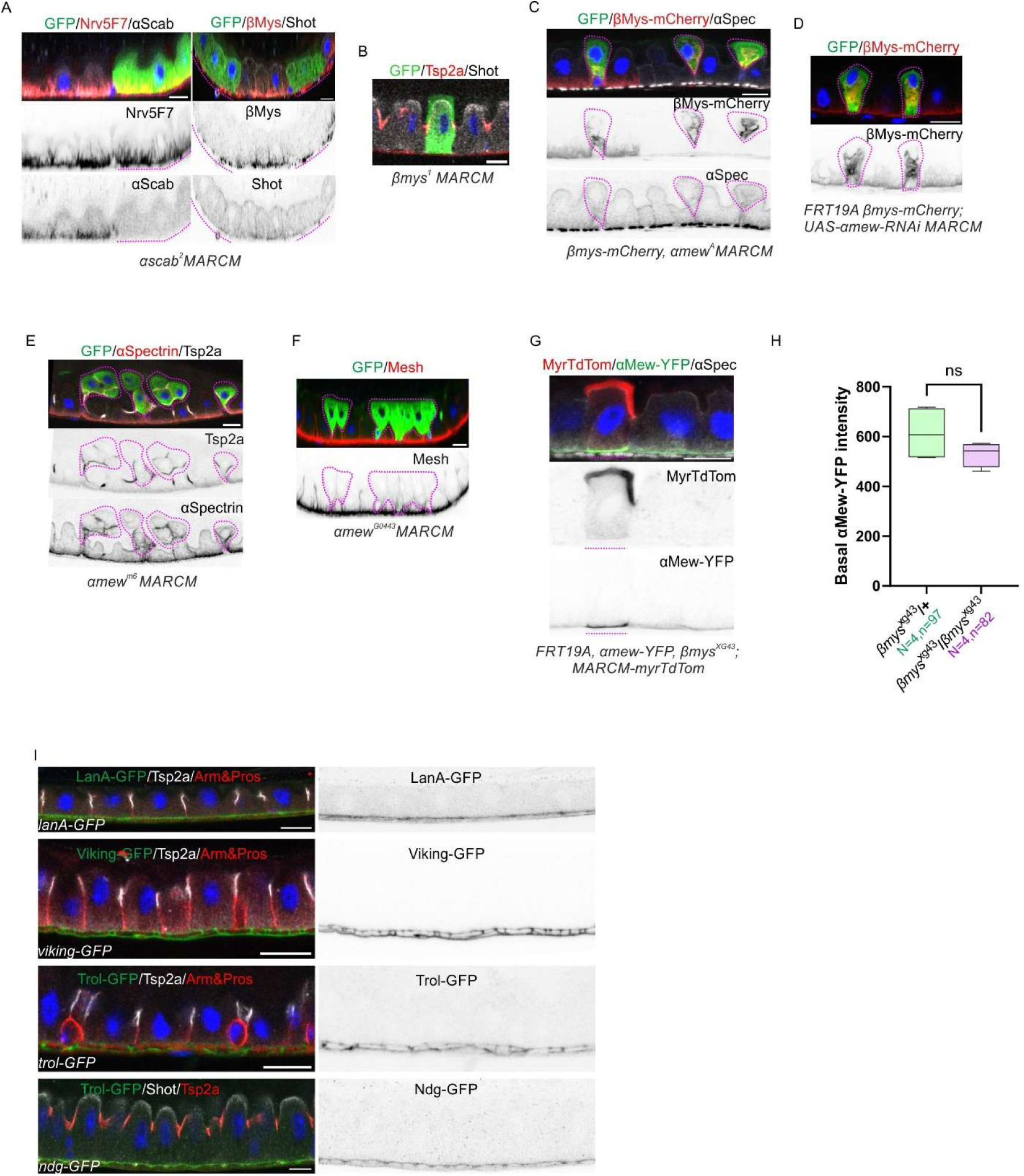
αMew/βν, not αScab/βMys dimers mediate the basal adhesion of enterocytes to ECM. A. Side views of *α scab*^2^ MARCM clones (GFP+, in green). The left panel shows no defect in the basal adhesion of the enterocytes or the structure of the basal labyrinth (BL), labeled with Nrv5f7 (red); the Scab antibody staining (grayscale) is lost in the clones; the right panel shows the loss of anti-βMys antibody staining in the *αscab*^2^ mutant clones. Single channels are shown below in inverted grayscale. The purple dotted line marks the mutant clones. B. A side view of a midgut containing a *βmys*^1^ MARCM clone (GFP+, green). The *βmys*^1^ mutant cell shows the same phenotype as *βmys*^XG43^ in Figure 3C, with no defect in basal adhesion. Tsp2a (SJ) in red and Shot in grayscale. C. A side view of a midgut containing *αmew*^A^ MARCM clones (GFP+, green) expressing βMys-mCherry. The mutant cells are apically extruded into the gut lumen. Both βMys-mCherry (red) and αSpectrin (grayscale) label the basal labyrinth. The βMys-mCherry and αSpectrin channels are shown below in inverted grayscale. The purple dotted line marks the mutant clones. D. A side view of a midgut containing MARCM clones (GFP+, green) expressing βMys-mCherry and *UAS-αmew-RNAi*. The cells in which αMew has been depleted are apically extruded into the gut lumen, a similar phenotype to the mutant clones in Figure 3E and panel C. βMys-mCherry (red, and as a single channel in inverted grayscale below) labels the basal labyrinth. The purple dotted line marks the mutant clones. E. A side view of a midgut containing *αmew*^m6^ MARCM clones (GFP+, green). The mutant cells are apically extruded into the gut lumen with well-formed septate junctions, labeled with Tsp2a (grayscale) and αSpectrin (red.) Individual channels of αSpectrin and Tsp2a are shown below in inverted grayscale. The purple dotted line marks the mutant clones. F. A side view of a midgut containing *αmew*^G0443^ MARCM clones (GFP+, green). The mutant cells are apically extruded into the gut lumen, but form normal septate junctions, labeled by Mesh (red, and as a single channel in inverted grayscale below). The purple dotted line marks the mutant clones. G. A side view of a midgut containing an *βmys*^XG43^ MARCM clone (Myr-TdTom+, red) expressing αMew-YFP. αMew localises basally in the *βmys*^XG43^ mutant enterocyte. Separate Myr-TdTom and αMew-YFP channels are shown in inverted grayscale below. The purple dotted line marks the mutant cell. H. Quantification of basal αMew-YFP signal intensity in *βmys*^XG43^ mutant ECs compared with the neighbouring wild type ECs, as shown in Figure S3G. Please note that as both *αmew-YFP* and *βmys*^XG43^ are on the X chromosome, there are 2 copies of *αmew-YFP* in the *βmys*^XG43^ homozygous mutant clones. The fluorescence intensity in the *βmys*^XG43^ mutant ECs was halved for comparison with the single copy of *αmew-YFP* in the heterozygous or WT neighbouring ECs. I. Side views of midgut epithelia expressing Laminin A, Viking (Collagen IV) Trol (Perlecan) and Nidogen endogenously-tagged with GFP. All four localise to the ECM on the basal side of adult midgut epithelium. LanA-GFP, Viking-GFP and Trol-GFP were stained with anti-GFP (green), and co-stained for Arm (AJ) and Pros (ee) in red and Tsp2a (SJ) in grayscale. Nidogen-GFP was stained with anti-GFP (green), Tsp2a (SJ; red) and Shot (apical domain; grayscale). Single GFP channels in inverted grayscale are shown on the right. Laminin and Nidogen were expressed from functional GFP-tagged transgenes under the control of their own promoters [16,19,20]. Viking and Trol were viable GFP protein trap lines[21]. Scale bars,10μm.

The only visible phenotype that we could detect was that in ⍺-*scab* mutant clones, βMys-mCherry failed to localise to the basal labyrinth and its levels were greatly reduced, consistent with the view that ⍺ and β Integrins must dimerise in the endoplasmic reticulum to be trafficked to the plasma membrane [18] (Figure 3A and Figure S3A). This indicates that βMys cannot dimerise with ⍺Mew in the enterocytes, as it does in ISCs and EBs, presumably because ⍺Mew preferentially pairs with βν. In support of this view, ⍺Mew does not re-localise to the basal labyrinth in *αscab* mutant cells (Figure 3B, Figure S3G and S3H). Thus, αMew cannot substitute for αScab in binding to βMys to induce its trafficking to the basal labyrinth.

### αMew/βν dimers mediate enterocyte adhesion to the basal ECM

In contrast to the lack of a phenotype in *αscab* and *βmys* mutant clones, enterocytes lacking either *αmew* or *βν* were extruded apically into the gut lumen (Figure 3E-G, and Figure S3C-F). This phenotype was stronger in *αmew* mutant ECs, which showed little to no basal attachment, whereas most *βν* mutant cells still retained some basal attachment (Figure 3F and 3G). When we examined the intensity of αMew at the basal domain in the absence of βν, a faint but detectable signal remained, which was ∼2.7-fold lower than in neighbouring wild-type enterocytes (Figure 3G and 3H). Thus, some αMew still localizes to the basal domain in *βν* mutant enterocytes, which may explain why αMew loss produces a stronger apical extrusion phenotype than loss of βν (Figure 3I).

In the adult midgut epithelium, the main ECM components, Laminin, Perlecan, Collagen IV and Nidogen, localize between the epithelium and the muscle layers (Figure S3I). Because αMew/βν is the primary integrin pair at the basal domain of the epithelial cells, it is likely to be the main mediator of adhesion to the ECM and downstream signaling.

### βMys can partially compensate for the absence of βν

The observations that *βν* mutant cells are only partially apically extruded from the midgut epithelium, whereas *αmew* mutant cells are completely extruded, and the persistence of a low level of basal αMew in *βν* mutant cells suggest that βMys can partially compensate for the loss of βν. We therefore examined the localisation of βMys in *βν* null mutant enterocytes. Confocal imaging of *βν*^SK4^ mutant clones showed that most of the βMys signal lies more apical than in the adjacent wildtype cells, suggesting that the basal labyrinth, where βMys localises, has become detached from the basal membrane (Figure 4A). This was confirmed by STED microscopy, which revealed that *βν* mutant cells showed a diffuse apical βMys signal above the basal domain, that co-localised with thread-like mCD8GFP membrane labelling (Figure 4B and 4C). Nevertheless, some βMys remained at the basal domain in the mutant cells (red arrowheads in Figure 4B and 4C).

**Figure 4.**
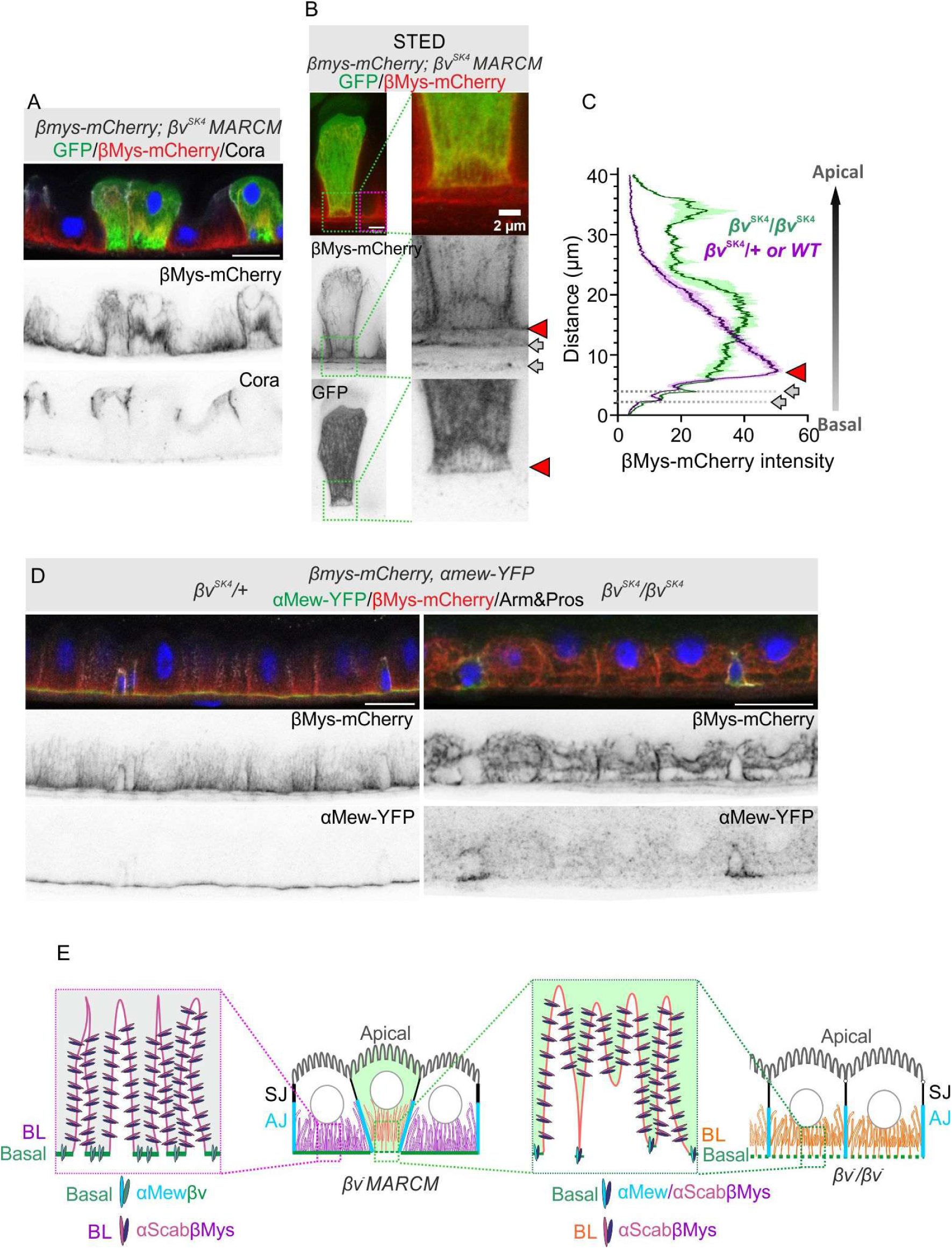
βMys can partially compensate for the absence of βν. A. A confocal image of a midgut expressing βMys-mCherry (red) and stained for Cora; (grayscale) containing *βν*^SK4^ MARCM clones (GFP+, green). Single channels of βMys-mCherry and Cora are shown below in inverted grayscale. Scale bar,10μm. B. A STED image of a *βν*^SK4^ MARCM clone stained as in (A), showing βMys-mCherry localization (red) in the mutant enterocytes. Single channels of βMys-mCherry and GFP are shown below in inverted grayscale. Scale bar, 5μm. The region outlined by the green dashed line is shown at higher magnification on the right. The purple dotted line marks a similar region in the neighbouring wild-type enterocyte. The red arrowhead marks the basal domain and the gray arrows indicate the positions of the muscle layers. Scale bar, 2μm. C. Quantification of the average intensities of βMys-mCherry fluorescence along the apical-basal axis in B. The purple line shows the data from the neighbouring WT enterocyte and the green line from the *βν*^SK4^ mutant enterocyte. D. A side view showing the localisation of αMew-YFP (green) and *β*Mys-mCherry (red) in *βν*^SK4^/+ and *βν*^SK4^/*βν*^SK4^ midgut epithelia. Arm (AJ) and Pros are in grayscale. The single αMew-YFP and βMys-mCherry channels are shown below in inverted grayscale. Scale bars, 10μm. E. Diagram showing the localisation of αMew/βν to the basal domain and of αScab/βMys to the basal labyrinth (BL) in WT enterocytes. In mutant enterocytes lacking *βν*, βMys can relocate to the basal domain and couple with αMew. The basal adhesion mediated by αMew/βMys is weaker than that mediated by αMew/βν in the neighbouring heterozygous enterocytes, and the mutant cells are therefore gradually outcompeted for contact with the basal ECM, leading to their extrusion into the gut lumen and the detachment of the basal labyrinth. In *βν*/*βν* homozygotes, all enterocytes have equally weak, basal adhesion mediated by αMew/βMys, and none of the cells are therefore extruded apically.

We exploited the fact that βν is non-essential to examine the midguts of *βν*/*βν* flies. Compared with *βν*/+ flies, βMys showed a similar, detached distribution in *βν*/*βν* midgut epithelia, with the detachment more pronounced at the centre of the cells than at the periphery (Figure 4D). The phenotype of both *βν* mutant clones and *βν*/*βν* homozygous epithelia suggests that most of the basal labyrinth has detached from ECM, leaving some attached fragments that are marked by thread-like mCD8-GFP signal and αSpectrin localization in the *βν* mutant clones (Figure 4B, Figure S4A and S4BB). In both cases, there is also a weak accumulation of βMys at the basal domain (Figure 4E). This suggests that in the absence of βν, βMys, the only remaining beta Integrin, compensates by localizing to the basal domain to maintain partial basal adhesion. βMys most probably forms heterodimers with *α*Mew basally, since about a third of the *α*Mew signal remains basal in *βν* mutant clones (Figure 3H). However, we cannot rule out the possibility that *α*Scab contributes to this weak adhesion to the ECM.

**Figure S4.**
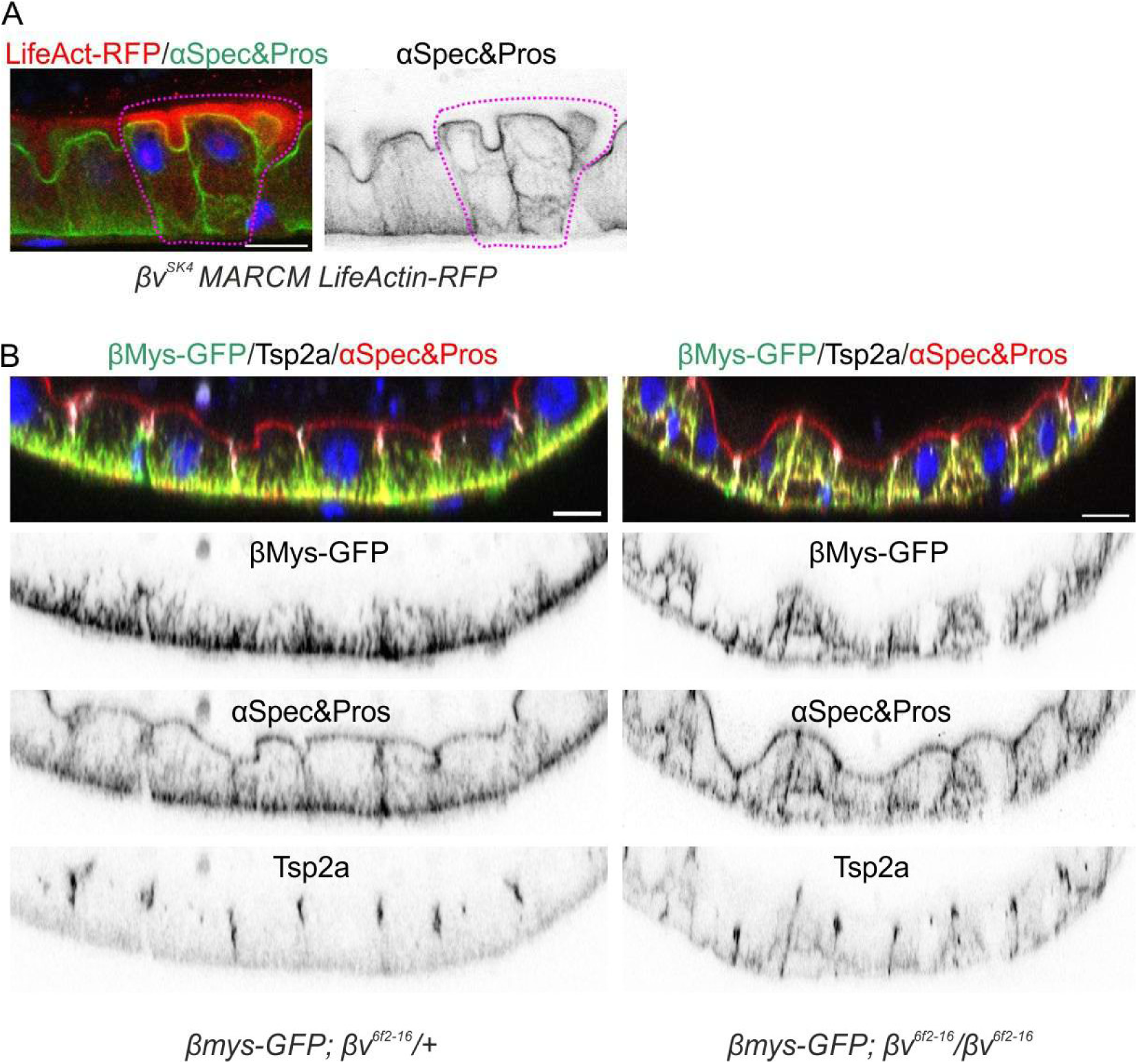
βMys can partially compensate for the absence of βν. A. A side view of a midgut containing a *βν*^SK4^ MARCM clone marked by LifeActin-RFP (red). The mutant cells have smaller basal domains in contact with the ECM, leading to their partial apical extrusion. The clone is outlined by the purple dotted line. αSpectrin (green, also shown as a single channel on the right in inverted grayscale) shows the detached basal labyrinth in the *βν*^SK4^ mutant enterocytes. B. Side views of βMys-GFP (green) in *βν*^6f2-16^/+ (left panel) and *βν*^6f2-16^/*βν*^6f2-16^ (right panel) midgut epithelia. The basal labyrinth, labeled by both βMys and αSpectrin (red), is detached from ECM in *βν*^6f2-16^/*βν*^6f2-16^. Pros (ee) also in red and Tsp2a (SJ) in grayscale. Single channels of αSpectrin, Pros and Tsp2a in inverted grayscale are shown below. Scale bars, 10μm.

### Wild-type enterocytes outcompete *βν* mutant enterocytes for adhesion to the basement membrane

Although *βν* mutant clones and the cells in *βν* homozygotes show a similar detachment of the basal labyrinth, they have different cell shapes. The *βν* mutant clones have smaller basal domains than the adjacent wild-type cells and are partially extruded, whereas the enterocytes in *βν* homozygotes have a normal cell shape and remain fully attached to the ECM (Figure 4A, 4B and Figure S4A versus Figure 4D and Figure S4B). Since the cells have the same genotype in each case, this difference is not cell-autonomous and must depend instead on the genotype of their neighbours. The simplest explanation for these observations is that mutant clones are outcompeted by their wild-type neighbours for adhesion to the basement membrane. The mutant cells only have low levels of αMew/βMys heterodimers, providing insufficient basal adhesion to resist competition from neighboring enterocytes, which use stronger αMew/βν– ECM interactions to occupy basement membrane space. By contrast, all enterocytes adhere equally weakly to the ECM in the *βν* homozygous epithelium, eliminating competition and preventing apical extrusion (Figure 4E).

### Enteroblasts out-compete *βν-/-* enterocytes for ECM adhesion and spread

Although the midguts of 14-28 day old *βν* homozygous adults have been reported to be disorganised and multilayered, with higher levels of ISC divisions (Okumura et al, 2014), the midguts of younger (4-7 days post eclosion) *βν* homozygotes appeared wild-type, forming a normal single-layered epithelium, with frequencies of dividing ISCs that were comparable to those of heterozygous controls (Figure 5A, Figure S5A and S5B). As mentioned above, αMew was strongly reduced at the basal sides of the enterocytes in the *βν/βν* epithelium (Figure 5A). However, the small, basal Pros-negative ISC/EBs retained high levels of αMew, presumably because these cells do not normally express *βν* and adhere to the ECM through αMew/βMys dimers (Figure 4D and Figure 5A). Some of these cells displayed a nearly triangular shape, in which their basal surfaces extend under the adjacent ECs, in contrast to the uniform morphology of basal progenitors in the *βν/+* epithelium (Figure S5C).

**Figure 5.**
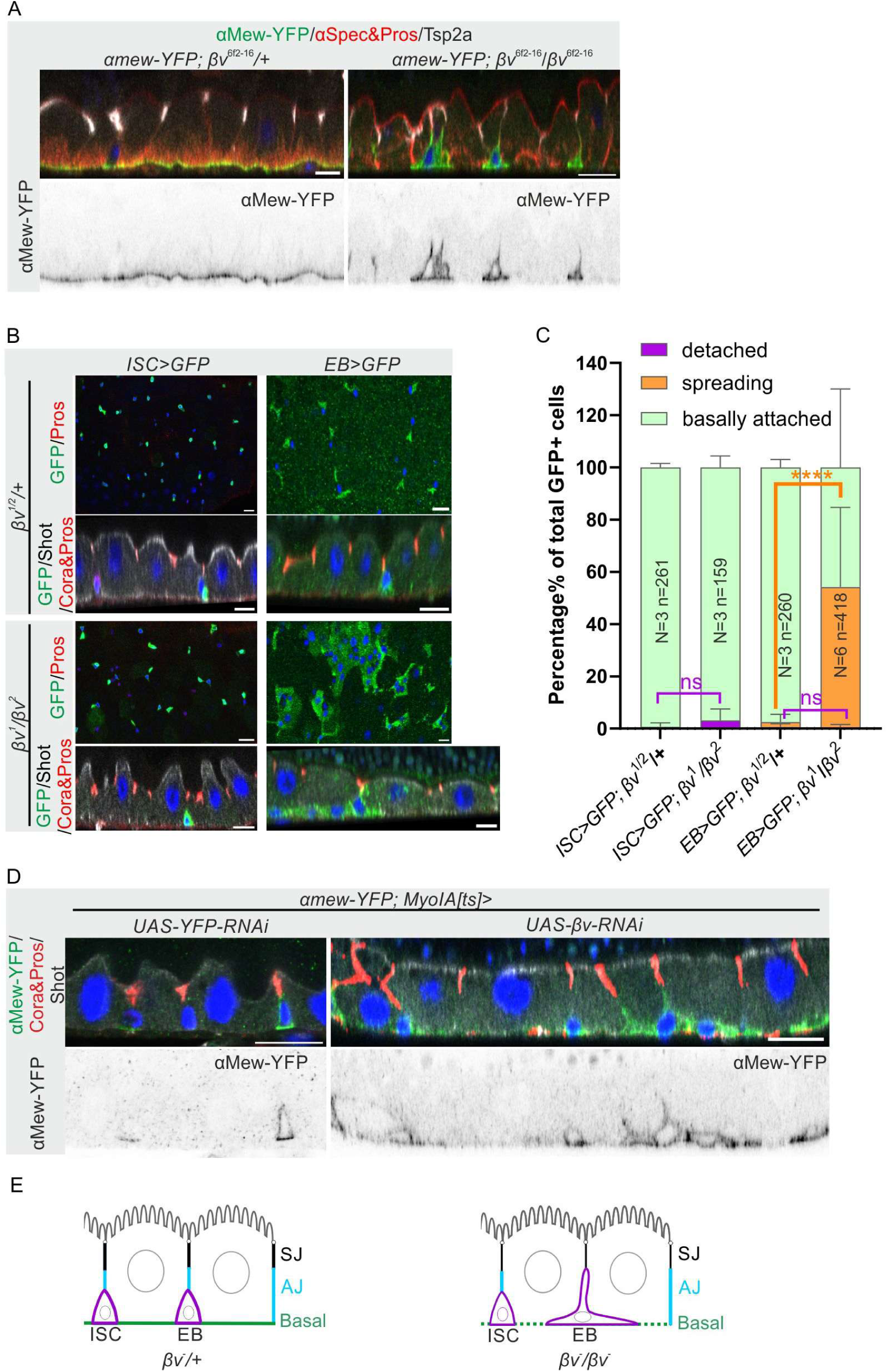
Enteroblasts spread basally in *βν* homozygotes. A. áMew-YFP localization in *βν/+* and *βν*/*βν* epithelia. The side views show that αMew-YFP (green) localises basally in *βν/+* enterocytes, but is much reduced in *βν*/*βν* enterocytes, and is only visible in the Pros-negative small cells, presumably ISCs/EBs. Some of the small cells have spread basally. αSpectrin and Pros in red, and Tsp2a in grayscale. The αMew-YFP channel is shown below in inverted grayscale. B. Basal surface and side views of *βν/+* and *βν*/*βν* epithelia, expressing either a GFP reporter for ISCs (*Delta-Gal4, UAS-mCD8GFP*, left) or EBs (*NRE>GFP*, right). Pros (red) labels the nuclei of the enteroendocrine cells; Cora (red) labels the SJs and Shot (grayscale) labels the apical domain. C. Quantification of the relative abundance of GFP+ ISCs or EBs with altered morphology in *βν/+* and *βν*/*βν* epithelia. Basally attached in light green, detached from ECM in purple and basal spreading in orange. N, number of experiments; n, number of cells. D. Side views of midgut epithelia expressing *αmew-YFP* (green) after knock-down of YFP (control) or *βν* specifically in the enterocytes using the *MyoIA[ts]* Gal4 driver. The EBs spread basally in the *MyoIA[ts]>βν-RNAi* midgut. Cora and Pros in red and Shot in grayscale. E. Diagram illustrating that ISCs and EBs are morphologically similar in *βν/+* heterozygous midguts, whereas the EBs spread basally in *βν*/*βν*. Scale bars, 10μm.

**Figure S5.**
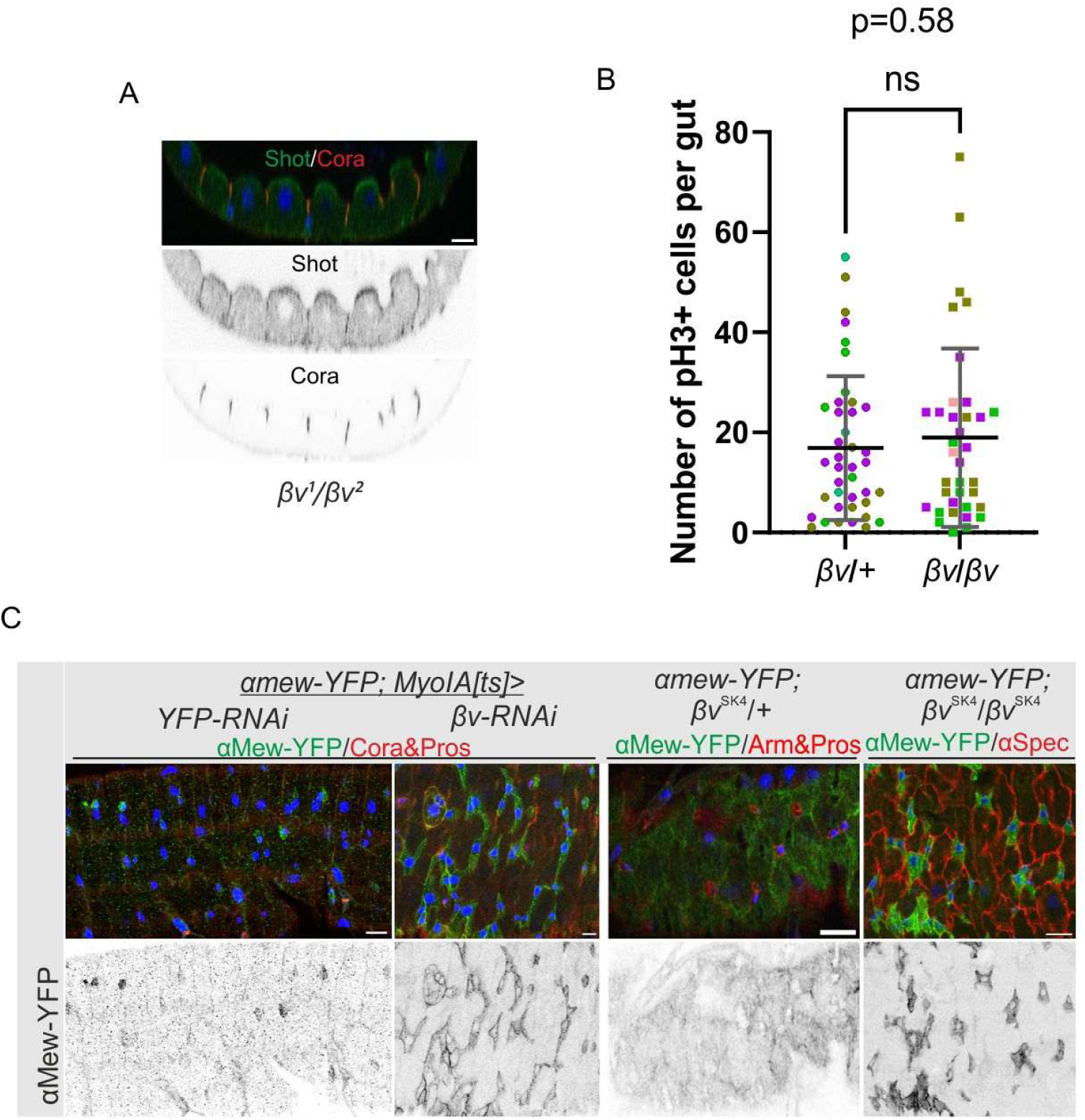
Enteroblasts spread basally in *βν* homozygotes. A. A side view of a *βν*^1^/*βν*^2^ midgut epithelium, showing a single layer of enterocytes with normal apical domains (Shot in green) and septate junctions (Cora in red). Single channels of Shot and Cora are shown below in inverted grayscale. B. Quantification of the number of phospho-histone 3 (pH3) positive cells in *βν/+* and *βν*/*βν* midgut epithelia. The ISC division rate was measured in *βν*^SK4^/*CyO* (green circles, Number of guts=11), *βν*^6f2-16^/*CyO* (purple circles, N=18) and *βν*^1^/*CyO* or *βν*^2^/*CyO* (brown circle, N =12); vs in *βν*^SK4^/ *βν*^SK4^ (green square, N=11), *βν*^6f2-16^/ *βν*^6f2-16^ (purple square, N=12) and *βν*^1^/*βν* ^2^ (brown square, N=14). The superplot was made using GraphPad Prism 9 software[22]. C. Basal surface views of the midgut epithelium as in Figure 5D & 5A, showing the basal expansion of the EBs when the enterocytes lack *βν*. Scale bars, 10μm.

To distinguish whether the small cells that spread basally in *βν* mutant midguts are ISCs, EBs or ees, we used *Delta-Gal4, UAS-mCD8GFP* as an ISC reporter, *NRE>GFP* as an enteroblast reporter and anti-Pros immunostaining for ee cells. Views of the basal surface and side views showed that only the EBs spread basally in *βν*/*βν* epithelia, compared to the *βν*/+ controls (Figure 5B and 5C).

Since depleting βν in just EBs does not cause them to spread basally (Figure 1D and 1E), we tested whether this phenotype was also due to a non-autonomous effect from the neighbouring enterocytes. Knocking down *βν* specifically in enterocytes using *MyoIA*^ts^ (Myo1a::Gal4, tub::Gal80^ts^) led to EB spreading, giving an identical phenotype to that seen in *βν* homozygous epithelia (Figure 5D and Figure S5C). Thus, the reduced adhesion of *βν* mutant enterocytes allows the EBs, which adhere to the basement membrane through αMew/βMys heterodimers, to outcompete them for contact with the ECM (Figure 5E).

The results above suggest that intestinal cells compete for adhesion to the ECM and squeeze out their neighbours if they are less adherent. To test this model, we asked whether reducing adhesion in the surrounding cells might also be able to rescue the stronger apical extrusion phenotype of *αmew* mutant ECs. We therefore generated *αmew* MARCM clones in a *βν* homozygous epithelium, so that the GFP+ cells lack both αMew and βν, while all other cells lack *βν* and only adhere weakly. Compared to the control *βν/+* epithelium, many fewer *αmew* mutant ECs detached from the basement membrane in the *βν* homozygous background (Figure 5A and 6C). Thus, the apical extrusion of *αmew* mutant ECs is also driven by the higher affinity of the neighbouring cells for the ECM, which allows them to outcompete the mutant cells for adhesion and thus displace them from the basement membrane. (Figure 5D)

**Figure 6.**
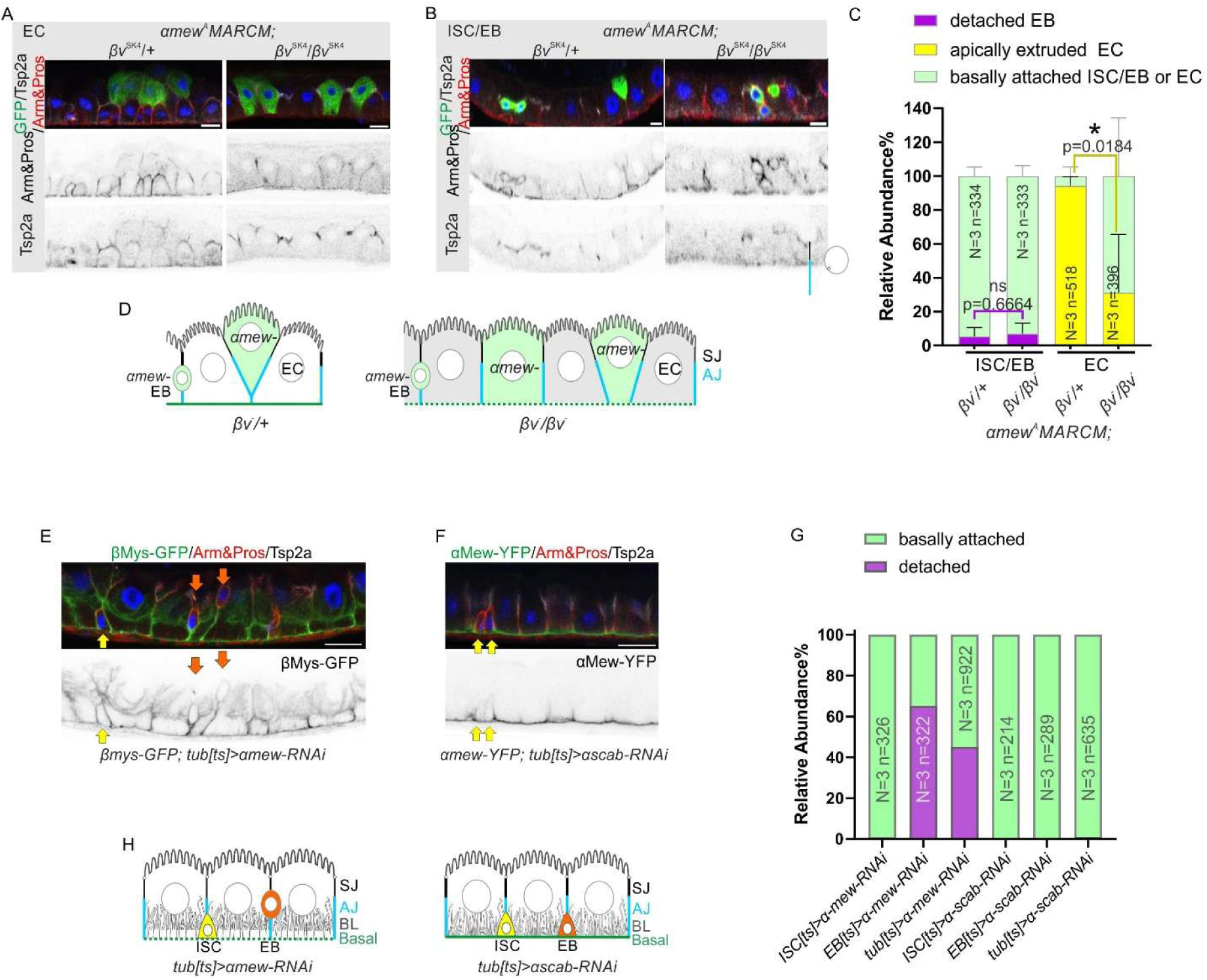
Basal adhesion mediated by αMew plays different roles in EBs and ECs. A. Side view of MARCM clones (GFP+, green) of *αmew*^A^ in the background of *βν*^SK4^/+ or *βν*^SK4^/*βν*^SK4^ midgut epithelium, showing that *αmew*^A^ mutant enterocytes have ameliorated apically extrusion phenotype in the background of *βν*^SK4^/*βν*^SK4^ comparing with *βν*^SK4^/+. B. Side views of *αmew*^A^ MARCM clones (GFP+, green) in *βν*^SK4^/+ or *βν*^SK4^/ *βν*^SK4^ epithelia, The Pros^-^ *αmew*^A^ mutant cells, presumably enteroblasts, show a similar detachment phenotype in both backgrounds. C. Quantification of the relative abundance of GFP+ ISC/EBs and ECs with different phenotypes: basally attached (green), detached but remain inside the epithelium (purple) and apically extruded into the gut lumen (yellow), in the experiments described in A&B: MARCM clones of *αmew*^A^ in the background of *βν*^SK4^/+ or *βν*^SK4^/ *βν*^SK4^ epithelia. D. A diagram illustrating how weakening overall basal adhesion by removing *βν* rescues the apical extrusion phenotype seen in *αmew*^A^ mutant enterocytes. However, removing βν does not rescue the detachment of *αmew*^A^ mutant enteroblasts. E. A side view of a midgut epithelium in which *αmew* has been knocked down in all cells using *tub*^ts^>*αmew-RNAi*. Some small Pros-negative cells have detached from the ECM and float inside the epithelium (orange arrows), while others have remained attached basally (yellow arrows). The enterocytes remain attached basally, but their basal labyrinths, marked by βMys-GFP have detached from the basal surface. Arm(AJ) and Pros in red, βMys-GFP in green and Tsp2a (SJ) in grayscale. The single βMys-GFP channel is shown below in inverted grayscale. F. A side view of midgut epithelium after tissue-wide knock down of *αscab* using *tub*^ts^>*αscab-RNAi*. The small Pros-cells remain attached basally (yellow arrows). Arm(AJ) and Pros in red, αMew-YFP in green and Tsp2a (SJ) in grayscale. The single αMew-YFP channel is shown below in inverted grayscale. G. A graph showing the relative abundance of basally-attached (light green) and detached ISCs/EBs (in purple). The *ISC*^ts^ and *EB*^ts^ > *αmew-RNAi* and *αscab-RNAi* experiments also included UAS-GFP to mark the cells subject to RNAi and the relative abundance is the percentage of GFP+ cells with the phenotype. In the *tub*^ts^ >*αmew-RNAi* or >*αscab-RNAi* experiments, the relative abundance is the percentage of small Pros-cells with the phenotype and does not distinguish between ISCs and EBs. N, total number of experiments; n, total number of cells. H. A diagram illustrating that losing αMew mediated adhesion in all cells in the epithelium using *tub^ts^>αmew-RNAi* have different effects on ISCs, EBs and ECs compared with control of *tub^ts^>αscab-RNAi*: In ECs, losing αMew at the tissue level does not cause detachment of the ECs but did cause the detachment of the basal labyrinth; ISCs (yellow) remain attached basally while EBs (orange) are detached. Scale bars, 10μm.

### The detachment of EBs lacking αMew may reflect a reattachment defect rather than reduced competition for basal adhesion

Although the detachment of *αmew* mutant ECs is partially rescued by the reduced adhesion of all ECs in *βν* homozygotes, this is not the case for the small cells in the epithelium, namely ISC/EBs, which show a similar frequency of detachment from the basement membrane in *βν* homozygotes compared to *βν/+* (Figure 5B and 6C). As shown in Figure 1, αMew is specifically required for the basal attachment of EBs, and not for ISCs, identifying the detached cells as EBs. This observation suggests that *αmew* mutant EBs do not become detached because they are outcompeted for adhesion to the basement membrane, but for some other reason (Figure 5D). To test this more rigorously, we removed all integrin-based adhesion by knocking down αMew in all cells in the epithelium using *tub^ts^>αmew-RNAi*. This did not cause detachment of the ECs, because all cells have similarly compromised basal adhesion, thereby removing competition, but did cause the detachment of the basal labyrinth, as previously observed in *βν* homozygotes (Figure 5E). However, the small Pros-cells consistently showed a detachment phenotype compared with the *αscab* RNAi controls (Figure 5E-G). Thus, loss of αMew in EBs causes a cell-autonomous detachment phenotype that is independent of the adhesion of their neighbours. Although the reason for this phenotype is unclear, one possibility is that it reflects a failure of the newborn EBs to re-attach to the basement membrane after an ISC division.

## Discussion

The *Drosophila* midgut epithelium is a homeostatic tissue that undergoes constant turnover, with the removal of damaged enterocytes and enteroendocrine cells and their replacement by differentiating EBs, which are then replenished by divisions of the ISCs. This homeostasis therefore requires tight coordination between the progenitor cells, their niche and their differentiated progeny. All these cell-types sit on the same basement membrane, but our results show that they interact with it in different ways using distinct combinations of α and β integrins (Figure 6).

**Figure 7.**
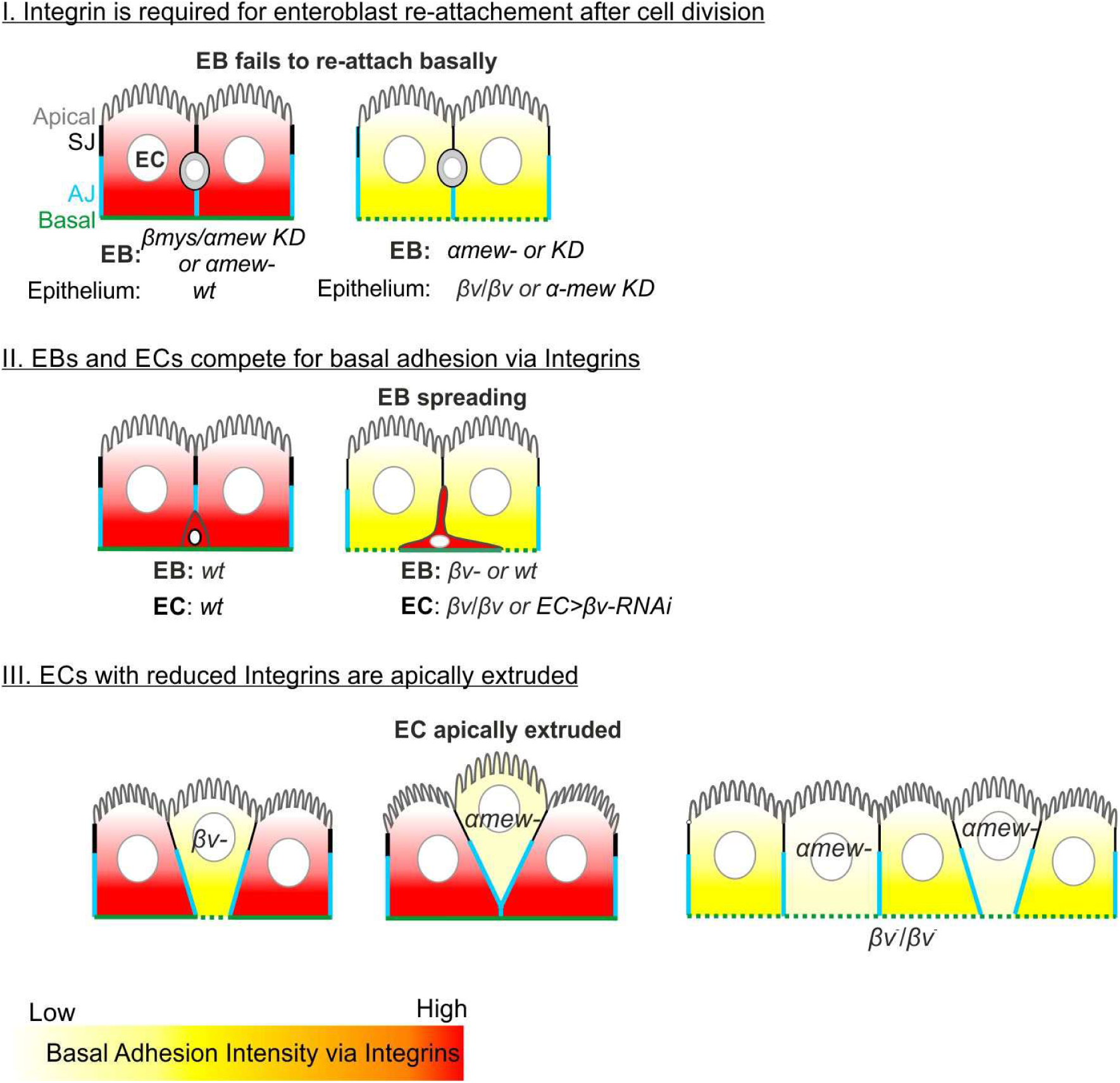
Cooperative and competitive contributions of Integrins to cell type-specific basal adhesion, EB reattachment post-division, and epithelial homeostasis.

The ISCs express αMew and βMys, but these mainly localise to the lateral sides of the cell, with only weak localisation to the basal membrane. Consistent with this, loss of either integrin has little effect on ISC adhesion to the basement membrane, since the ISCs rarely detach, although mutant ISCs divide less frequently and are gradually lost over time [9,11]. Thus, integrin signalling promotes ISC growth and survival, but attachment to the ECM is mediated by other ECM receptors, possibly functioning in parallel to the integrins. This may be related to the observation that the adhesion of ISCs to the basement membrane depends on their cell-autonomous secretion of the heparan-sulphate proteoglycan, Perlecan [23]. This makes Dystroglycan, which binds Perlecan, a good candidate for an ECM receptor that functions redundantly with Integrins to mediate ISC basal attachment [24].

Unlike ISCs, EBs require αMew and βMys for their basal attachment, since up to 60% of EBs are found above the basement membrane in the middle of the epithelium when αMew, βMys or Talin are knocked down in these cells. In contrast to ECs, this loss of contact with the ECM is not influenced by the genotype of the neighbouring cells, indicating that it is not a result of competition for adhesion to the ECM. Two reports have proposed that the asymmetric ISC divisions that give rise to EBs are oriented at an oblique angle to the plane of the epithelium, whereas the symmetric divisions lie in the plane [10,25]. If this is the case, EBs are born above the plane of the epithelium and must extend basally to reattach to the ECM and achieve their normal position. This suggests that the cell-autonomous detachment phenotype of EBs lacking αMew or βMys arises because these integrin dimers function to reattach apically-positioned EBs to the ECM after asymmetric ISC divisions. This reattachment must be rapid, however, because imaging dividing ISCs at 1 frame/ hour did not detect newborn EBs above the basement membrane [26].

Unlike ISCs and EBs, which only express significant amounts of αMew and βMys, enterocytes also express αScab and βv. This triggers a reassortment of the integrins with αMew pairing with βv rather than with βMys. This presumably occurs because βv binds more strongly to αMew than βMys does, leaving the latter to pair with αScab. Consistent with this, knockdown of αScab leads to the disappearance βMys rather than its association with αMew. This suggests that βMys is degraded if it cannot bind αScab in the endoplasmic reticulum to form dimers that can be transported to the plasma membrane.

The enterocytes are the only *Drosophila* cells that are known to express two β integrins, but this occurs in several mammalian cell-types. For example, keratinocytes express both β1 and β4 integrins, with the former localising to dynamic focal adhesions and the latter forming stable hemidesmosomes [27]. The localisations of βMys and βv in midgut enterocytes are even more strikingly different, with αMew/βv dimers localising to the basal surface in contact with the basement membrane and αScab/βMys dimers localising to the basal labyrinth. These non-overlapping distributions could arise because the two integrins take different trafficking routes to the plasma membrane or because they are retained in each region by binding to different extracellular ligands. αMew is the orthologue of mammalian α6 integrin, which binds to Laminin, consistent with the idea that αMew/βv binds to the ECM to attach ECs to the basement membrane. It is less clear what αScab/βMys bind in the basal labyrinth, which lacks obvious electron-dense ECM material in electron micrographs [17]. However, the septate junction protein, Mesh, which localises to the basal labyrinth, is the orthologue of the newly-identified ECM component and Integrin ligand, SNED1, making it a potential partner for αScab/βMys [28]. It is also unclear whether αScab/βMys serve any function at the basal labyrinth, as the latter’s structure is unchanged in the mutants. One possibility is that this provides a mechanism to sequester βMys away from the basal surface so that αMew/βv can link the ECs to the basement membrane.

Unlike αScab/βMys, αMew/βv dimers function as the main ECM receptors that anchor the enterocytes to the basement membrane, since cells lacking either integrin detach and are eventually extruded. The phenotype of *βv* mutant clones is weaker than that of *αmew* clones, suggesting that βMys can partially compensate. This may be because a small amount of βMys can form dimers with αMew rather than αScab in the absence of βv, or because some of the αMew/βMys dimers that attach EBs to the basement membrane perdure into the ECs. More significantly, the detachment of integrin mutant enterocytes depends on the genotype of their neighbours. *βv* mutant clones detach when surrounded by heterozygous cells, but all ECs remain attached to the basement membrane in *βv* homozygotes, which are viable. Similarly, the ECs in *αmew* mutant clones detach when surrounded by *αmew/+* cells, but remain attached if all ECs lack βv. This reveals that the enterocytes compete with each other for attachment to the basement membrane, with the less adherent cells being extruded. It is therefore tempting to suggest that down-regulation of αMew/βv may act as a trigger to delaminate dying ECs and shed them into the gut lumen.

Although competition between epithelial cells for adhesion to the basement membrane has not previously been demonstrated, it is likely to be a common phenomenon. For example, the Rac-dependent down-regulation of α6/β4 Integrin in intestinal crypt cells reduces the area of their basal surface area in contact with the basement membrane, leading to a cell shape change that bends the epithelium at the crypt/villus boundary to form the hinge [29]. In several contexts, reducing integrin adhesion in single cells, leads to their detachment from the basement membrane and apical extrusion. Talin1 knockdown in isolated cells in mammary acini in 3d culture triggers their apical extrusion into the lumen [30]. Similarly, loss of β4 Integrin in isolated basal keratinocytes leads to their delamination from the basal layer and differentiation, and the Notch signal that induces differentiation down-regulates β4, suggesting this may trigger delamination from the basal layer [31,32]. Furthermore, this mechanism has been co-opted to remove unfit epidermal stem cells, with DNA damage activating Notch to down-regulate integrins and induce cell extrusion and differentiation [33].

Not only do enterocytes compete for adhesion to the basement membrane, but they also compete with the EBs, since the EBs expand their basal surface area when the enterocytes lack βv. This raises the question of why EBs and ECs express different β Integrins and whether this is impacts gut development. One attractive scenario is that βMys and βv fulfil similar roles to β1 and β4 integrins in keratinocytes, with the former forming transient attachments that facilitate cell migration and the latter forming stable attachments in hemidesmosomes [27]. EBs are migratory cells that can move to sites of damage, and αMew/βMys dimers may form dynamic ECM attachments that aid these movements, like β1 Integrin in keratinocytes [34,35]. By contrast, αMew/βv dimers are likely to form stable ECM attachments that anchor the enterocytes in place, fulfilling a similar role to β4 Integrin. This raises the possibility that changes in integrin levels and competition between EBs and ECs may regulate EB migration. For example, the extrusion and shedding of damaged ECs into the gut lumen is likely to involve the down-regulation of αMew/βv to detach the cells from the basement membrane, and this could promote the movement of EBs to this site through competition for ECM adhesion, where they can differentiate to replace the dying EC.

## Materials and methods

### Drosophila melanogaster stocks

*y*^2^ or attp2 (BDSC #25710) flies were used as wild type unless otherwise specified. Other stocks used in this study were as follows:

<u>Fluorescently tagged protein lines:</u>

αMew-YFP[36] (Kyoto DGGR#115524), αScab-GFP (BDSC#63170), αInflated-YFP[36] (Kyoto DGGR#115467), βMys-GFP[37] (gift from Nick Brown, University of Cambridge), Mys-mcherry[38] (gift from Nick Brown, University of Cambridge), βν-GFP[16] (VDRC#v318193), LanA-GFP[16] (VDRC#v318155); Viking-GFP[21] (BDSC#98343), Trol-YFP[36] (Kyoto DGGR#115613), Nidogen-GFP[39] (BDSC#66766).

<u>Mutant stocks:</u>

FRT19A *αmew*^A^ (Kyoto DGGR#117029), FRT18A *αmew*^m6^ (BDSC#1483), FRT19A *αmew*^G0443^ (Kyoto DGGR#111926), FRT42D *αscab*^2^ (BDSC#68157), FRT19A *βmys*^XG43^ (gift from Nick Brown, University of Cambridge), FRT19A *βmys*^1^ (BDSC#23862), *βν*^1^ (gift from Nick Brown, University of Cambridge), *βν*^2^ (BDSC#600168), FRT40A *βν*^SK4^ (NIG-fly# M2L-0662), *βν*^6f2-16^ (this study)

<u>UAS Gal4 responder lines:</u>

EB^ts^ (Su(H)GBE^ts^): Su(H)GBE-Gal4, UAS-mCD8GFP; tub>Gal80^ts^, UAS-GFP

ISC^ts^(Delta^ts^): tub>Gal80^ts^; Delta-Gal4, UAS-mCD8GFP

tub^ts^: tub>Gal80^ts^; tub-Gal4 (BDSC#86328)

EC^ts^ (MyoIA^ts^): MyoIA-Gal4, tub>Gal80^ts^

<u>EB/ISC GFP reporter lines:</u>

βν^1^/CyO; Delta-Gal4, UAS-mCD8GFP

βν^2^/CyO; Delta-Gal4, UAS-mCD8GFP

βν^1^/CyO; NRE>GFP (BDSC#30728)

βν^2^/CyO; NRE>GFP (BDSC#30728)

<u>UAS-RNAi lines</u>[40]:

UAS-αmew-RNAi (BDSC#44553), UAS-αscab-RNAi (BDSC#38959), UAS-βmys-RNAi (BDSC#33642), UAS-βν-RNAi (BDSC#61916), UAS-talin-RNAi (BDSC#33913).

<u>The following stocks were used to generate (positively labelled) MARCM clones</u> [41]:

MARCM FRT19A GFP: w, hsFLP, tubP-GAL80, FRT19A;; tubP-GAL4, UAS-mCD8::GFP/TM3, Sb

MARCM FRT19A myr-TdTomato: w, hsFLP, tubP-GAL80, FRT19A;; tubP-GAL4, UAS-myr-TdTomato /TM3, Sb

MARCM FRT19A βν: w, hsFLP, tubP-GAL80, FRT19A; FRT40A *βν*^SK4^; tubP-GAL4, UAS-mCD8::GFP/TM3, Sb

MARCM FRT40A GFP: hsFLP[1]; FRT 40A tubP-GAL80; tubP-GAL4, UAS-mCD8::GFP (gift from Nick Brown, University of Cambridge)

MARCM FRT40A LifeAct-RFP: hsFLP[1]; FRT 40A tubP-GAL80; tubP-GAL4, UAS-LifeAct-RFP (gift from Nick Brown, University of Cambridge)

MARCM FRT42D GFP: hsFLP[1], Act-Gal4, UAS-mCD8::GFP; FRT 42D tubP-GAL80 (this study)

### Stock maintenance

Standard procedures were used for *Drosophila* husbandry and experiments. Flies were reared on standard fly food supplemented with live yeast at 25 °C. For the conditional expression of UAS-RNAi in adult flies, parental flies were crossed at 18 °C and the resulting offspring reared at the same temperature until eclosion. Adult offspring were collected for 3 days and then transferred to 29 °C to inactivate the temperature sensitive GAL80^ts^ protein. To generate MARCM clones, flies were crossed at 25 °C and the resulting offspring were subjected to heat shocks either as larvae (from L2 until eclosion) or as adults (5–9 days after eclosion). Heat shocks were performed at 37 °C for 1 h twice daily. Flies were transferred to fresh food vials every 2–3 days and kept at 25 °C for at least 4 days after the last heat shock to ensure that all wild-type gene products from the heterozygous progenitor cells had turned over. All samples used in this study were obtained from adult female flies.

### Heat Fixation

The heat fixation protocol is based on a heat–methanol fixation method used for *Drosophila* embryos[42]. Detailed procedure was described previously[5,43]. Samples were dissected in PBS, transferred to a wire mesh basket, and fixed in hot 1X TSS buffer (0.03% Triton X-100, 4 g/L NaCl; 95 °C) for 3 s before being transferred to ice-cold 1X TSS buffer and chilled for at least 1 min. Subsequently, samples were transferred to washing buffer and processed for immunofluorescence staining.

### Antibody stainings

Antibody stainings were performed as described previously[5,43]. After blocking in blocking buffer (1xPBS, 0.1%TritonX-100, 5%NGS (vol/vol)), samples were incubated with the appropriate primary antibody/antibodies diluted in blocking buffer at 4 °C overnight. Following several washes in washing buffer (1xPBS, 0.1%TritonX-100), samples were incubated with the appropriate secondary antibody/antibodies either at room temperature for 2 h or at 4 °C overnight. Samples were then washed several times in washing buffer and mounted in Vectashield containing DAPI (Vector Laboratories) on borosilicate glass slides (No. 1.5, VWR International). All antibodies used in this study were tested for specificity using clonal analysis (MARCM) or RNAi.

#### Primary antibodies

Mouse monoclonal antibodies: anti-Cora (C566.9) (RRID:AB_1161642), anti-αSpec (3A9) (RRID:AB_528473), anti-Arm (N2 7A1) (RRID:AB_528089), anti-Pros (MR1A) (RRID:AB_528440), anti-Nrv1 (Nrv5f7)(RRID:AB_528395) and anti-βMys (CF.6G11) (RRID:AB_528310). The monoclonal antibodies against Cora (developed by Fehon, R., University of Chicago), αSpec (developed by Branton, D. / Dubreuil, R., Harvard University), Arm (developed by Wieschaus, E., Princeton University), Pros (developed by Doe, C.Q., University of Oregon), Nrv1 (developed by Salvaterra, P.M., City of Hope, Beckman Research Institute) and βMys (developed by Brower, D., Harvard Medical School) were obtained from the Developmental Studies Hybridoma Bank, created by the NICHD of the NIH and maintained at The University of Iowa, Department of Biology, Iowa City, IA 52242, and used at 1:100 dilution.

Polyclonal antibodies: Rabbit anti-RFP (Cat#MBL-PM005, Medical & Biological Laboratories (MBL), 1:500 dilution) (RRID:AB_591279); Rabbit anti-Mesh (RRID:AB_2568117) and anti-Tsp2A (gift from Mikio Furuse, National Institute for Physiological Sciences, Okazaki, Japan, 1:1,000 dilution); Rat anti-βν and anti-Scab (gift from Nakanishi Yoshinobu, Kanazawa University, Japan, anti-βν lot 53 (corresponding to aa 753-799 in the intracellular region), 1:100 dilution; anti-Scab lot 63 (corresponding to aa 235-284 in the extracellular region), 1:300 dilution); Rabbit anti-PH3 (Millipore Cat# DAM1545035, 1:500 dilution) (RRID:AB_2315134); Chicken anti-GFP (Abcam, Cat. #ab13970, 1:1,000 dilution) (RRID:AB_300798); Guinea pig anti-Shot (gift from Nashchekin D. University of Cambridge, 1:1,000 dilution).

#### Secondary antibodies

Alexa Fluor secondary antibodies (Invitrogen) were used at a dilution of 1:1,000.

Alexa Fluor 488 goat anti-mouse (#A11029), Alexa Fluor 488 goat anti-rabbit (#A11034), Alexa Fluor 488 goat anti-guinea pig (#A11073), Alexa Fluor 488 goat anti-chicken IgY (#A11039), Alexa Fluor 555 goat anti-mouse (#A21422), Alexa Fluor 555 goat anti-rabbit (#A21428), Alexa Fluor 568 goat anti-guinea pig (#A11075), Alexa Fluor 647 goat anti-mouse (#A21236), Alexa Fluor 647 goat anti-rabbit (#A21245). Only cross-adsorbed secondary antibodies were used in this study to eliminate the risk of cross-reactivity.

For STED imaging, following secondary antibodies were used at dilution of 1:500.

ChromoTek GFP-Booster ATTO647N (Chromotek, Cat. #gba647n) (RRID:AB_2629215), ChromoTek RFP-Booster ATTO594 (Chromotek, Cat. #rba594), abberior STAR RED goat anti-rabbit IgG (Abberior, STRED-1002-500UG) (RRID:AB_2631390), abberior STAR ORANGE goat anti-chicken IgY (Abberior, STORANGE-1005-500UG) (RRID:AB_2857377), FluoTag-X4 anti-GFP abberior STAR 635p (NanoTag Biotechnologies Cat# N0304-AB635P-L) (RRID:AB_3075902), Anti-rabbit IgG Atto 594 (goat polyclonal) (Sigma Aldrich, Cat. #77671) (RRID:AB_1137663).

### Immunofluorescence Imaging

Confocal images were collected on an Olympus IX81 (40×1.35 NA Oil UPlanSApo, 60× 1.35 NA Oil UPlanSApo) using the Olympus FluoView software Version 3.1 and processed with Fiji (ImageJ).

### Stimulated emission depletion (STED) super-resolution imaging

The microscope design is a variant of a STED system described in detail previously [44,45]. STED imaging was performed on a custom built system centred around a 775 nm depletion laser (Katana HP, OneFive) with an 80 MHz repetition rate and pulse length of 700 ps. Pulsed excitation wavelengths at 488 nm (PicoQuant) 590 nm (SuperK Extreme, NKT Photonics) and 656 nm (PicoQuant) were available. The system used an 8 kHz resonance mirror for beam scanning, custom electronics for pulse timing, and avalanche photodiodes with three detection channels centered around 525 nm, 625 nm and 705 nm. Data acquisition and instrument control was performed using a custom control program developed in the LabView environment (National Instruments). STED images were taken with 590 and 656 nm excitation, 19 or 24 nm pixel size, 2048 x 2048 image format and 650 accumulation per line. STED depletion laser power was 120 mW at the microscope base camera side port. The detailed settings of the STED microscope are described previously[46].

### Generation of *βν*^6f2-16^ flies and characterisation of *βν*^SK4^ flies

We used the CRISPR/Cas9 method[47] to generate a null allele of *βν*. sgRNA was in vitro transcribed from a DNA template created by PCR from two partially complementary primers:

forward primer: 5′-GAAATTAATACGACTCACTATA<u>ttgcgagcccccggattacgtgg</u>GTTTTAGAGCTAGAAATAGC-3′; reverse primer: 5′-AAAAGCACCGACTCGGTGCCACTTTTTCAAGTTGATAACGGACTAGCCTTATTTTAACTTGCTATTTCTA GCTCTAAAAC-3′.

The sgRNA was injected into *Act5c-Cas9* embryos[48]. Putative *βν* mutants in the progeny of the injected embryos were recovered, balanced, and sequenced. The *βν*^6f2-16^ allele contains a 2bp deletion around the CRISPR site, resulting in one missense mutation and a frameshift that leads to amino acid change from 725 and a stop codon after amino acid 749, covering the region corresponding to the transmembrane domain (727-749), which is shared by both isoforms.

The *βν*^SK4^ allele is originated from NIG-Fly KO collection (described in FBrf0254534), this allele contain 28bp deletion creating frameshift which leads to amino acid change from 146 and a stop codon after aa 167, the resulting peptide only contain less than a third of the extracellular head region of βν, possibly also deleting the vWA domain (βA domain) which is shared by both isoforms and important for ligand binding.

Since the antibody against βν was supplied with limited quantity, we compared phenotypes of *βν*^6f2-16^*/βν*^6f2-16^ and *βν*^SK4^*/βν*^SK4^, *βν*^6f2-16^ or *βν*^SK4^ trans-heterozygous with other known *βν* null alleles to *βν* null alleles homozygous flies, found it displayed the same phenotype in midgut epithelium as in *βν*^1^*/βν*^1^, *βν*^1^*/βν*^2^ and *βν*^2^*/βν*^2^, thus we concluded both the *βν*^6f2-16^ and *βν*^SK4^ alleles are also loss of function alleles.

### Measuring αMew-YFP, βMys-mcherry fluorescence intensities in wild type enterocytes and Mys-mcherry fluorescence intensity in *βν* mutant enterocytes

We used the straight-line tool (width=2μm, length=15μm in wild type EC and 40 μm in *βν* mutant EC) in Fiji to mark the regions from basal to apical in enterocytes. 20 lines in 3 images of αMew-YFP, βMys-mcherry fluorescence intensities in wild type enterocytes, and 14 lines in 3 images of βMys-mcherry fluorescence intensity in *βν* mutant enterocytes obtained on STED microscope were measured to plot the average fluorescence intensity. Graph was made by GraphPad Prism 9 software. Representative images and graphs were shown in Figure 2A and 2B, Figure S2E, Figure 5B and 5C.

### Measuring αMew-YFP fluorescence intensities at the basal domain in *βmys*^XG43^ and *βν*^SK4^ mutant enterocytes and their neighbouring wild type cells

We used the straight-line tool (width=2μm, length to cover the basal domain of individual enterocyte) in Fiji to measure the αMew-YFP fluorescence intensity in *βν*^SK4^ and *βmys*^XG43^ mutant enterocytes and their neighbouring wild type cells as control. Graph was made by GraphPad Prism 9 software. Representative images and graphs were shown in Figure 3G and 3H, and Figure S3G and S3H.

## Data availability

All fly stocks and reagents used in this study are available on request.

## Acknowledgements

We would like to thank Gurdon Institute Imaging Facility for microscopy and image analysis support, and members of the St Johnston laboratory for their advice, discussion and support. We are very grateful to Mihoko Tame, Dmitry Nashchekin, Tony Ip, Mark Peifer, Thomas Lecuit, Stefan Luschnig, Benjamin Klapholz, Nicholas Brown, Golnar Kolahgar, Mikio Furuse, Graham Thomas, Denise Montell, Fernando Díaz-Benjumea, Benoit Biteau, Akira Nakamura, Dorothea Godt, the Bloomington *Drosophila* stock centre, the NIG, the Kyoto *Drosophila* Stock Center and the Developmental Studies Hybridoma Bank for providing fly stocks and antibodies.

The authors declare no competing financial interests.

## Author contributions

J. Chen and D. St Johnston conceived and designed the project, prepared the figures and wrote and edited the manuscript. The STED imaging experiments on wild type and *βν* mutant was performed by D. Sugita and E. Allgeyer; the experiment on *βν-RNAi MARCM* clones was performed by D. Saumya; the experiment on αMew’s localisation in *βν*/*βν* midgut was performed D. Shunmugam; and the rest were performed by J. Chen. The project funding, administration, and supervision were provided by D. St Johnston.

